# Natural regeneration in tree islands drives long-term recovery of native forest-associated woody species in oil palm landscapes

**DOI:** 10.64898/2026.11.07.765027

**Authors:** Vivien Michel, Fabian Brambach, Nathaly Guerrero-Ramírez, Dian Muh Fauzan, Bambang Irawan, Leti Sundawati, Dirk Hölscher, Holger Kreft, Gustavo Brant Paterno

**Affiliations:** Biodiversity, Macroecology and Biogeography, University of Göttingen, Göttingen, Germany; Forest Botanical Garden and Arboretum, University of Göttingen, Göttingen, Germany; Centre of Biodiversity and Sustainable Land Use (CBL), University of Göttingen, Göttingen, Germany; Flora Division, Celebica Conservation Action Foundation (Yayasan Aksi Konservasi Celebica), Palopo, Indonesia; Forestry Department, Faculty of Agriculture, University of Jambi, Jambi, Indonesia; Department of Forest Management, IPB University, Kampus IPB Darmaga, Bogor, Indonesia; Tropical Silviculture and Forest Ecology, University of Göttingen, Göttingen, Germany; Land Use Transformation Systems (PUI-PT BLasTS) Centre of Excellence, University of Jambi, Jambi, Indonesia

**Keywords:** ecological restoration, enrichment planting, Indonesia, natural succession, non-native species, sustainable palm oil, tree diversity experiment, TreeDivNet

## Abstract

1. Large-scale conversion of tropical rainforests to oil palm plantations is a major driver of biodiversity loss in Southeast Asia, particularly threatening native forest-associated plant species and their associated biota. Integrating tree islands within monocultural oil palm landscapes has emerged as a promising approach to mitigate biodiversity loss. However, their effectiveness in sustaining natural regeneration and long-term recovery of native forest-associated plant species remains unclear.
2. We analyzed temporal trajectories of diversity, abundance, and composition of naturally recruiting woody species in tree islands embedded in a monoculture-dominated landscape. We used a large-scale biodiversity enrichment experiment with 52 tree islands, varying in area (25 - 1600 m²) and planted tree diversity (0, 1, 2, 3, or 6 species), established in a conventional oil palm plantation in Sumatra, Indonesia, in 2013.
3. Across four censuses conducted between 2 and 8 years after the onset of natural regeneration, we found 82 species of recruiting woody plants (trees and shrubs) belonging to 64 genera and 34 families. At the landscape (gamma) scale, species richness and abundance increased by ∼42% and ∼69%, respectively, from 2018 to 2024, with shifts in community evenness and continued species arrival. After 8 years, neither richness nor abundance saturated, indicating continued accumulation of recruiting plant species.
4. Tree islands promoted natural regeneration of native diversity, with 90% of species and individuals being native. Native forest-associated species (N = 35) increased steadily at the landscape scale, while localized dominance of alien species did not indicate arrested succession.
5. At the local (alpha) scale, larger island area exerted a sustained positive effect on recruiting diversity. Higher initial planted tree diversity promoted native recruitment, particularly of forest-associated species, at earlier stages, an effect that diminished over time.
6. *Synthesis and applications.* Tree islands promote natural regeneration and accelerate recovery of native forest-associated diversity over extended periods, providing an effective strategy to enhance forest-associated biodiversity within oil palm landscapes. Preserving the last remaining lowland rainforests, irreplaceable reservoirs of native forest-associated species, remains essential to support continued recruitment into tree islands that can create stepping-stone habitats and a more wildlife-friendly agricultural matrix.

## 1 Introduction

Southeast Asian lowland rainforests are centers of exceptional biodiversity and endemism (Cai et al., 2023; Myers et al., 2000; Pillay et al., 2022). Yet, these ecosystems have experienced rapid deforestation and substantial biodiversity loss in recent decades, largely driven by agricultural expansion, especially large-scale oil palm monocultures (Gaveau et al., 2022; Ke et al., 2025; Qaim et al., 2020; Rembold et al., 2017). At the landscape scale, these changes have also promoted the spread of alien plant species, further homogenizing and simplifying species-rich communities (Kabache et al., 2026; Rembold et al., 2017). With continued forest cover loss and fragmentation, recovering native forest-associated diversity is an increasing priority (Gaveau et al., 2022; Mungi et al., 2021; Mungi et al., 2025). Alongside efforts to safeguard remaining forests (Gibson et al., 2011; Watson et al., 2018), there is a critical need to understand the temporal dynamics of the recovery of native biodiversity within ecologically simplified oil palm landscapes (Brancalion et al., 2025; Dislich et al., 2016).

A continuum of ecological restoration approaches used to enhance native biodiversity in human-modified tropical landscapes (Chazdon et al., 2021) ranges from natural regeneration to more intensive interventions such as tree enrichment and large-scale tree planting. Natural regeneration can produce species-rich secondary forests (Metz et al., 2026; Norden et al., 2009; Rozendaal et al., 2019; Van Breugel et al., 2013), but in fragmented landscapes this is often constrained by limited seed availability and disperser scarcity, particularly for later-successional species (Bardino et al., 2023; Chazdon & Guariguata, 2016; Norden et al., 2009; Poorter et al., 2023; Zahawi & Augspurger, 2006). Large-scale tree planting can accelerate succession by rapidly creating shaded, favourable microclimatic conditions for later-successional species (Brancalion et al., 2025; Holl et al., 2020; Paterno et al., 2024; Poorter et al., 2023). Tree islands - clusters of native trees planted within agricultural or degraded landscapes - offer an intermediate strategy by acting as regeneration nuclei through improved seed dispersal and microclimatic conditions (Brancalion et al., 2025; Holl et al., 2020; Paterno et al., 2024; Poorter et al., 2023; Song et al., 2025; Zahawi et al., 2013;), and may increase connectivity as stepping-stone habitats (Arroyo-Rodríguez et al., 2020; Saura et al., 2014; Uezu et al., 2008). Compared with full plantings, tree islands and natural regeneration deliver comparable restoration outcomes in terms of biodiversity and ecosystem functioning (Banin et al., 2022; Crouzeilles et al., 2017; Joyce et al., 2025), while requiring lower costs and offering greater scalability (Chazdon & Guariguata, 2016; Jones et al., 2018).

Natural regeneration is a long-term process shaped by interacting factors that govern community assembly, including restoration design factors such as the diversity and species identity of planted trees or the area of tree islands (Brancalion et al., 2025; Paterno et al., 2024; Piquer-Doblas et al., 2024). Larger areas can support more species through higher colonization and lower extinction rates, as predicted by the species–area relationship (Arrhenius, 1921) and the equilibrium theory of island biogeography (MacArthur & Wilson, 1967). However, most evidence remains short-term, highlighting the need to assess restoration trajectories over longer times frames (Brancalion et al., 2025; Meli et al., 2017), particularly the temporal dynamics of how tree island design (i.e., planted tree diversity and island area) influences biodiversity recovery at both local and landscape scales.

Taxonomic diversity in restored forests typically increases over time (Rozendaal et al., 2019; Vellend et al., 2013), but alien species can suppress native species regeneration over extended periods, a process termed arrested succession (Matos et al., 2022; Royo & Carson, 2006; Vellend et al., 2013). The rapid establishment, fast growth, and persistence of alien species in disturbed open habitats (D’Antonio & Meyerson, 2002; Gioria et al., 2023; Kabache et al., 2026; Poorter et al., 2023) allow dominant alien species to alter community structure and ecosystem processes, substantially constraining recovery of native diversity (D’Antonio & Meyerson, 2002; Pearson et al., 2018; Weidlich et al., 2020). Understanding how restoration design shapes the long-term trajectories of native and alien diversity is therefore essential.

Open canopies, hot, dry microclimates, and intensive management in oil palm landscapes, including weeding, fertilization, and herbicide application, exclude most forest-dependent plant species, driving declines in native plant diversity and cascading losses of forest-dependent fauna (Foster et al., 2011; N. Guerrero-Ramírez, unpublished data; Rembold et al., 2017;). Promoting native forest-associated species is therefore a central objective of tropical restoration (e.g., Cole et al., 2010; Gilman et al., 2016; Lu et al., 2017; Paterno et al., 2024; Simões et al., 2024). A key challenge is identifying restoration strategies capable of restoring plant diversity in oil palm landscapes, providing refugia for native forest-associated species while limiting invasion by alien plants. Addressing this knowledge gap requires long-term studies of temporal restoration trajectories (Mungi et al., 2025; Weidlich et al., 2020).

Here, we analyzed temporal changes in the taxonomic diversity and abundance of native (including forest-associated) and alien recruiting plant species, covering 8 years of natural regeneration dynamics with four surveys within the EFForTS-BEE experiment (Ecological and Socioeconomic Functions of Tropical Lowland Rainforest Transformation Systems: Biodiversity Enrichment Experiment; Teuscher et al., 2016). This large-scale biodiversity enrichment experiment comprises 52 tree islands established in an industrial-scale oil palm plantation in Sumatra in 2013, with free succession starting in 2014 (Teuscher et al., 2016). The tree islands systematically vary in area (25, 100, 400, 1600 m²) and planted tree diversity (0, 1, 2, 3, or 6 native species; Fig. 1).

**Figure 1.**
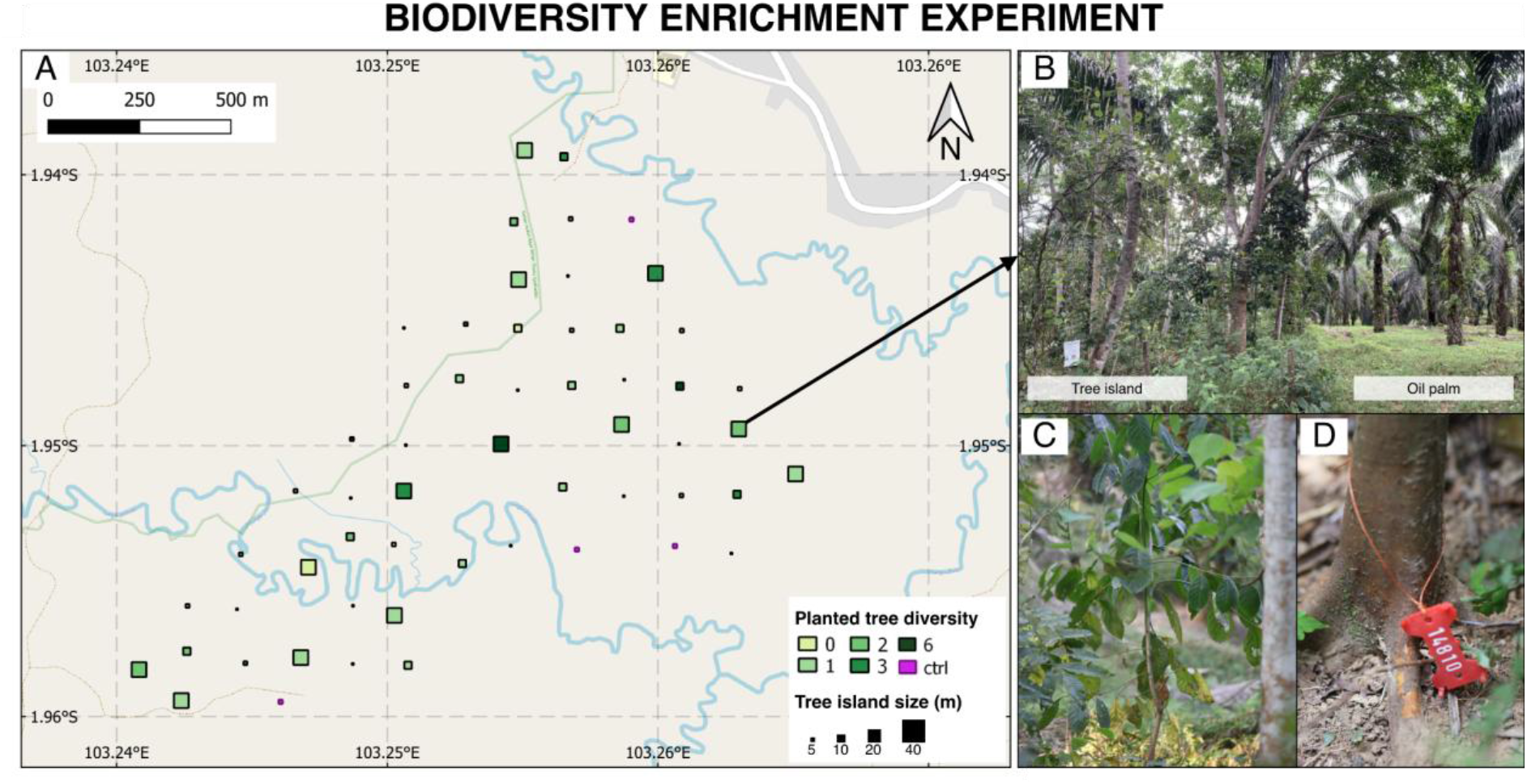
Overview of the EFForTS-BEE study system in Jambi, Sumatra, Indonesia. **(A)** Spatial arrangement of experimental tree islands embedded within an industrial oil palm plantation, showing variation in tree island area and planted tree diversity. **(B)** Example of a tree island within the surrounding oil palm matrix. **(C)** Naturally recruiting woody vegetation within a tree island. **(D)** Example of an individually tagged naturally recruiting woody plant. All naturally recruited woody individuals > 130 cm in height were surveyed and assigned unique tag identifiers. Photos: B, Gustavo Paterno (2025); C, Tika Simanjuntak (2024); D, Vivien Michel (2024).

This study addresses two main research questions: **(1)** How do the diversity, abundance and composition of naturally recruiting species change over time, and do tree islands promote native forest-associated species while limiting alien dominance at the landscape (gamma) scale? and **(2)** How do tree island area and planted tree diversity influence temporal changes in local (alpha) recruiting diversity, particularly of native forest-associated species?

At the landscape scale, we expect overall recruiting diversity and abundance to increase over time (Brancalion et al., 2025; Rozendaal et al., 2019; Vellend et al., 2013) (H1), driven by cumulative seed arrival and vegetation development (Poorter et al., 2023). A continuous increase in diversity is expected if long-distance seed dispersal by animals (Piquer-Doblas et al., 2024) is an important mechanism in the study region; saturation may occur if the regional species pool is limited (Poorter et al., 2023); and a slight decline could result from competitive exclusion (Royo & Carson, 2006), such as suppression of native recruitment by dominant alien species (D’Antonio & Meyerson, 2002; Matos et al., 2022; Pearson et al., 2018; Vellend et al., 2013). Native forest-associated recruiting diversity is expected to show a delayed but sustained increase (H2), as conditions become increasingly shaded over time and suitable for later-successional species (Poorter et al., 2023; Van Breugel et al., 2013). In contrast, alien recruiting diversity is expected to saturate earlier (H3), reflecting rapid early colonization abilities but a smaller regional species pool (D’Antonio & Meyerson, 2002; Gioria et al., 2023; Poorter et al., 2023).

At the local scale, we hypothesize that larger tree islands support greater recruiting diversity, including native forest-associated diversity, with effects strengthening over time (H4), owing to the species-area relationship (Arrhenius, 1921; MacArthur & Wilson, 1967; Rosenzweig, 2003), reduced edge effects, and greater microhabitat heterogeneity promoting disturbance-sensitive later-successional species (Poorter et al., 2023; Van Breugel et al., 2013). Higher planted tree diversity is expected to initially increase recruiting diversity, particularly native forest-associated diversity, by enhancing canopy closure and structural and environmental heterogeneity facilitating contrasting ecological strategies among recruiting species (Poorter et al., 2023; Van der Sande et al., 2024). However, this effect is expected to weaken over time as natural recruitment increases vegetation structural complexity (Kikuchi et al., 2024) and reduces differences in canopy and microclimatic conditions among tree islands (H5). An overview of all hypotheses is provided in the supplementary materials (Table S1).

## 2 Materials and methods

### 2.1 Study region

The study was conducted in the EFForTS-BEE experiment located near Bungku village, Jambi, Sumatra, Indonesia (01.95° S, 103.25° E, 47 ± 11 m a.s.l.). The region was originally dominated by dipterocarp lowland rainforest, which had been converted into conventional oil palm plantations (Laumonier et al., 2010; Teuscher et al., 2016). The surrounding landscape is now dominated by oil palm plantations (83%), with smaller areas of fallow land (6%), bare soil (5%), secondary forest (4%) and rubber monocultures (2%), as well as orchards, water bodies and urban infrastructure (<1%) (Khokthong, 2019). Scattered trees, i.e., trees outside forests, occur at a density of 10.1 trees per ha (Korol et al., 2021). The linear distance to the nearest forest, Hutan Harapan, is approximately 18 km, estimated using Google Earth (Google, 2025). The climate is humid tropical, with a mean temperature of 26.7 °C and an annual rainfall of 2235 mm, marked by two rainfall peaks around March and December and a drier period during July to August (Drescher et al., 2016). Loamy Acrisol soils are the predominant soil type in the region (Allen et al., 2015).

### 2.2 Experimental design

The EFForTS-BEE experiment was established within a 140-ha industrial oil palm plantation of PT. Humusindo Makmur Sejatwe (Teuscher et al., 2016; Zemp et al., 2023; Fig. 1). Oil palms had been planted in a 9 x 9 m triangular grid between 2001 and approximately 2007, with ca. 143 oil palms per ha (Teuscher et al., 2016). Conventional plantation management includes fertilizer application, manual weeding and removal of epiphytes, with rare application of herbicides (Teuscher et al., 2016). The plantation is also used for livestock farming (Teuscher et al., 2016). A total of 52 tree islands were established within the oil palm plantation in December 2013, systematically varying in area (25, 100, 400, 1600 m²) and levels of planted tree diversity (0, 1, 2, 3, or 6 species), following a random partition design (Bell et al., 2009). Natural regeneration was allowed within the tree islands (Teuscher et al., 2016; Zemp et al., 2023). The minimum distance between tree islands is 85 m.

Within each tree island, native trees were planted in a 2 m grid among oil palms. To improve light availability for planted trees, approximately 40% of oil palms were removed, except in the 5 x 5 m islands that were positioned between existing oil palm rows. Six native multi-purpose species were used: *Parkia speciosa* Hassk. (Fabaceae), *Archidendron jiringa* (Jack) I.C.Nielsen (Fabaceae) and *Durio zibethinus* L. (Malvaceae) that are mainly grown for fruits, *Peronema canescens* Jack (Lamiaceae) and *Rubroshorea leprosula* (Miq.) P.S.Ashton & J.Heck (Dipterocarpaceae) used for timber, and *Dyera polyphylla* (Miq.) Steenis (Apocynaceae) that produces natural latex (Paterno et al., 2024; Teuscher et al., 2016). After planting, no fertilizers, herbicides, or pesticides were applied within the tree islands anymore, and after May 2016, 2.5 years after establishment, manual weed control was restricted to 1 m circles around the planted trees to promote natural regeneration (Zemp et al., 2023). The EFForTS-BEE experiment is part of the global network of tree diversity experiments (TreeDivNet; Paquette et al., 2018). Further methodological details are provided in Teuscher et al. (2016).

### 2.3 Data collection

All experimental plots were fully censused for naturally regenerating woody plants from 2 to 8 years after the onset of natural regeneration in May 2016. Censuses were conducted in 2018 (April - August), 2020 (October - December), 2022 (June - October), and 2024 (August - November).

The survey included trees, shrubs, and bamboo with a minimum height of 130 cm. In 2024, lianas were also surveyed (Table S2) but were not included in our analyses. Recruiting oil palms were included if they had a minimum stem diameter of 10 cm. All recorded individuals were tagged and, when still alive, resurveyed across census years. Species identification was carried out to species level, with a few exceptions (*Secamone* sp., *Syzygium* sp., *Ixora* sp. and *Nephelium* sp.), which were identified to genus level. Identification was based on field photographs and voucher specimens, supported by relevant floras (Flora Malesiana, Tree Flora of Malaya, Flora of the Malay Peninsula) and databases (Plants of the World Online, Digital Flora of Indonesia), as well as taxonomic monographs and revisions where necessary. For each species, lifeform (tree, treelet, shrub, liana) and habitat (forest-associated, open-habitat-associated) were assigned using the same data sources. Species were classified as native or alien to Sundaland using Plants of the World Online (2025). For each recorded individual, height, stem diameter at 10 cm (d10), and stem diameter at 130 cm (dbh) were measured.

### 2.4 Data analysis

Prior to analysis, data from the four separate censuses were cleaned and standardized, allowing for accurate temporal comparisons of recruiting diversity. For more information on data processing, please refer to the supplementary materials.

To estimate diversity at the local tree island (alpha) and landscape (gamma) scales, we used the Hill numbers framework (Chao et al., 2021). In this framework, q = 0 corresponds to species richness (^0^*D*), q = 1 reflects the effective number of common species (^1^*D,* exponential of Shannon entropy), and q = 2 describes the effective number of dominant or highly abundant species (^2^*D*, inverse of the Simpson concentration index) (Chao et al., 2021).

Hill diversity was calculated for ^0^*D*, ^1^*D,* and ^2^*D* using the *iNEXT.3D* package (Chao et al., 2021). Taxonomic diversity was estimated for five subsets: (i) all species, (ii) native species, (iii) native forest-associated species, (iv) native open-habitat-associated species, and (v) alien species, at both landscape and local scales for each survey year (2018, 2020, 2022, 2024). Abundance was calculated as the number of individuals, and basal area was calculated from the stem diameters at 10 cm.

Pielou’s evenness was calculated with the *diversity* function in the R package *vegan* (Oksanen et al., 2025). Community composition was analyzed using principal coordinates analysis (PCoA) based on Bray–Curtis dissimilarities calculated from Hellinger-transformed species abundance data using the R packages *vegan* (Oksanen et al., 2025) and *ape* (Paradis & Schliep, 2019). Prior to analysis, seven plots were excluded because they lacked recruiting individuals in at least one census year (*N*_5 × 5 m_ = 6 and *N*_10 × 10 m_ = 1). Statistical differences in species composition among years were tested using permutational multivariate analysis of variance (PERMANOVA) using the function *adonis2* in *vegan* (Oksanen et al., 2025), with plot identity included as a strata term to account for repeated measurements through time. Multivariate dispersion was assessed using the *betadisper* function in *vegan* (Oksanen et al., 2025).

To test for the experimental effects of tree island area and planted tree diversity on local recruiting diversity over time, we fitted linear mixed models (LMMs) for the three focal subsets of native, native forest-associated, and alien species at the local scale. The two continuous variables tree island area and planted tree diversity, plus year as categorical variable, were included as fixed effects. Interaction terms between year and planted tree diversity, and between year and tree island area, were added to test for time-dependent effects on recruiting diversity. Tree island ID was included as a random effect to account for repeated measurements. Models were fitted using the *glmmTMB* function from the R package *glmmTMB* (Brooks et al., 2017). Fixed effects significance was evaluated with the Wald Chi-square test (type III sum of squares), using the *Anova* function in the R package *car* (Fox & Weisberg, 2019). Model predictions were visualized with the *ggpredict* function from R package *ggeffects* (Lüdecke, 2018). The response variable, recruiting diversity, was log-transformed (ln(x+1)) to improve model residuals. The predictor variables were scaled and log-transformed as ln(x) for tree island area and ln(x+1) for planted tree diversity. For model validation and residual diagnostics, we used the *check_model* function from the R package *performance* (Lüdecke et al., 2021). For all statistical tests, a significance level of *P*-value < 0.05 was used.

All analyses were performed in R version 4.4.1 (R Core Team, 2024). Data cleaning and calculations were performed with the R package *tidyverse* (Wickham et al., 2019). Figures were created using the R packages *ggplot2* (Wickham, 2016) and *patchwork* (Pedersen, 2025), next to *ggeffects* (Lüdecke, 2018) and *sjPlot* (Lüdecke, 2025).

## 3 Results

### 3.1 Temporal changes in recruiting diversity and abundance at the landscape scale

Our dataset comprised 12,829 records of naturally regenerated woody individuals, collected across four censuses (2018, 2020, 2022, 2024), including resurveyed individuals. Across surveys, we found a total of 82 woody species (Table S3, excluding lianas) among 8,237 identified individuals that established within the tree islands, representing 64 genera from 34 plant families.

The great majority of recruiting species were native (90%, *N* = 74), as were 90% of recorded individuals. Thirty-seven species (46%) were forest-associated, 35 of which were native. The majority of alien species were associated with open habitats (*N* = 6), only two were forest-associated (*Swietenia macrophylla* King (Meliaceae), *Artocarpus heterophyllus* Lam. (Moraceae)). The most species-rich families throughout the years were Moraceae (*N* = 12), Euphorbiaceae (*N* = 10), and Rubiaceae (*N* = 8). The number of recruiting individuals and the number of species, genera, and families increased over time, while basal area increased to 6.04 m² ha^-1^ by 2024 (Fig. 2; Fig. S1).

**Figure 2.**
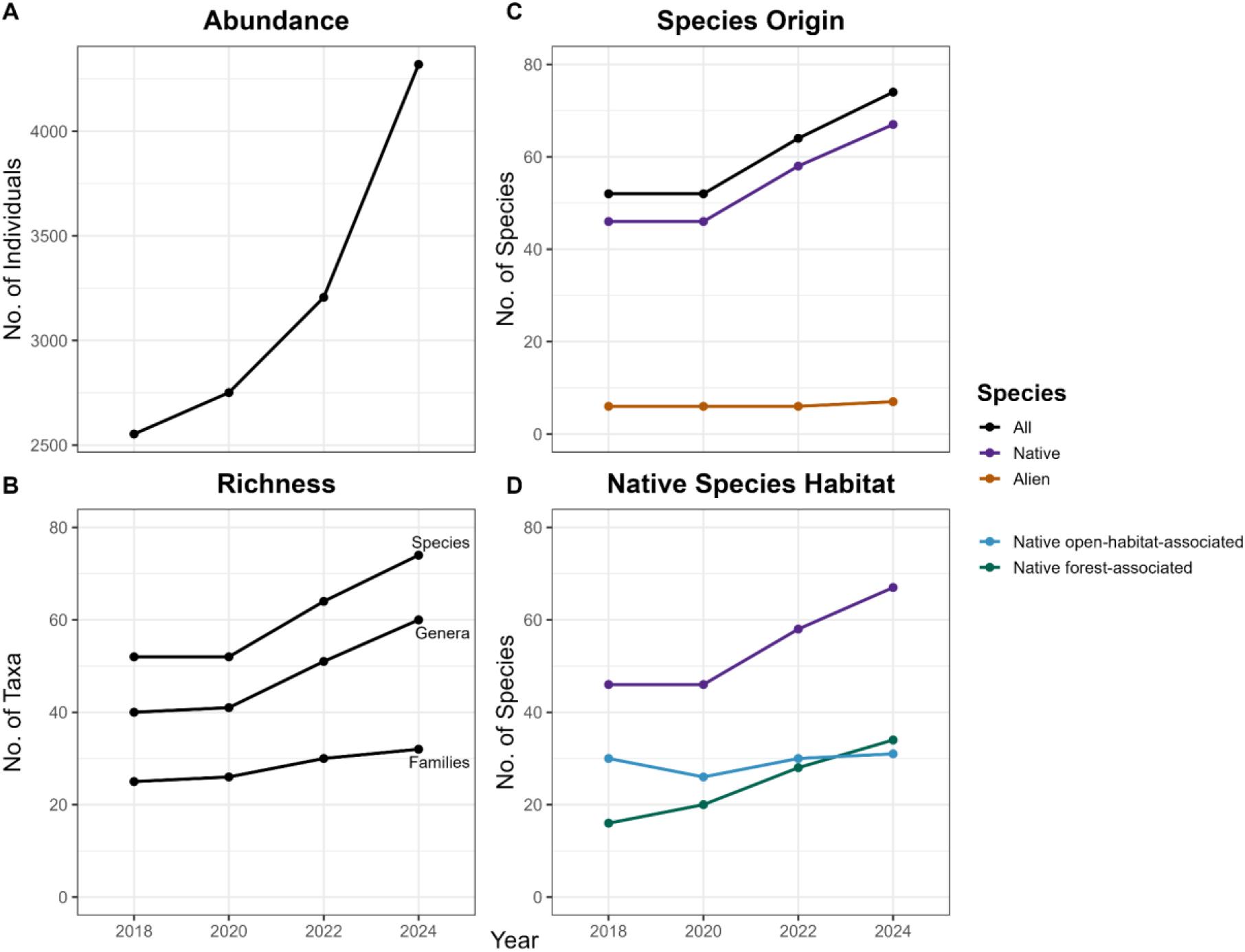
Temporal changes in naturally recruiting woody vegetation within the EFForTS-BEE tree islands from 2018 to 2024. Panels show changes in **(A)** total abundance of individuals, **(B)** taxonomic diversity at species, genera, and family levels, **(C)** numbers of native and alien species (biogeographic status), and **(D)** habitat associations of native species (forest-associated and open-habitat-associated). Values represent cumulative diversity and abundance across all tree islands at the landscape (gamma) scale.

Between 2018 and 2024, we documented a 42% increase in species richness at landscape scale (^0^*D*_2018_ = 52, ^0^*D*_2024_ = 74), while seven more families appeared. Recruiting abundance increased by 69% (*N*_2018_ = 2553, *N*_2024_ = 4319). For both species richness (^0^*D*) and Shannon diversity (^1^*D*), rarefaction curves showed increases over time (Fig. 3A; Fig. S2A). For Simpson diversity (^2^*D*), there was no further increase after 2022 (^2^*D*_2022_ = 8.59, ^2^*D*_2024_ = 7.64; Fig. S2A).

**Figure 3.**
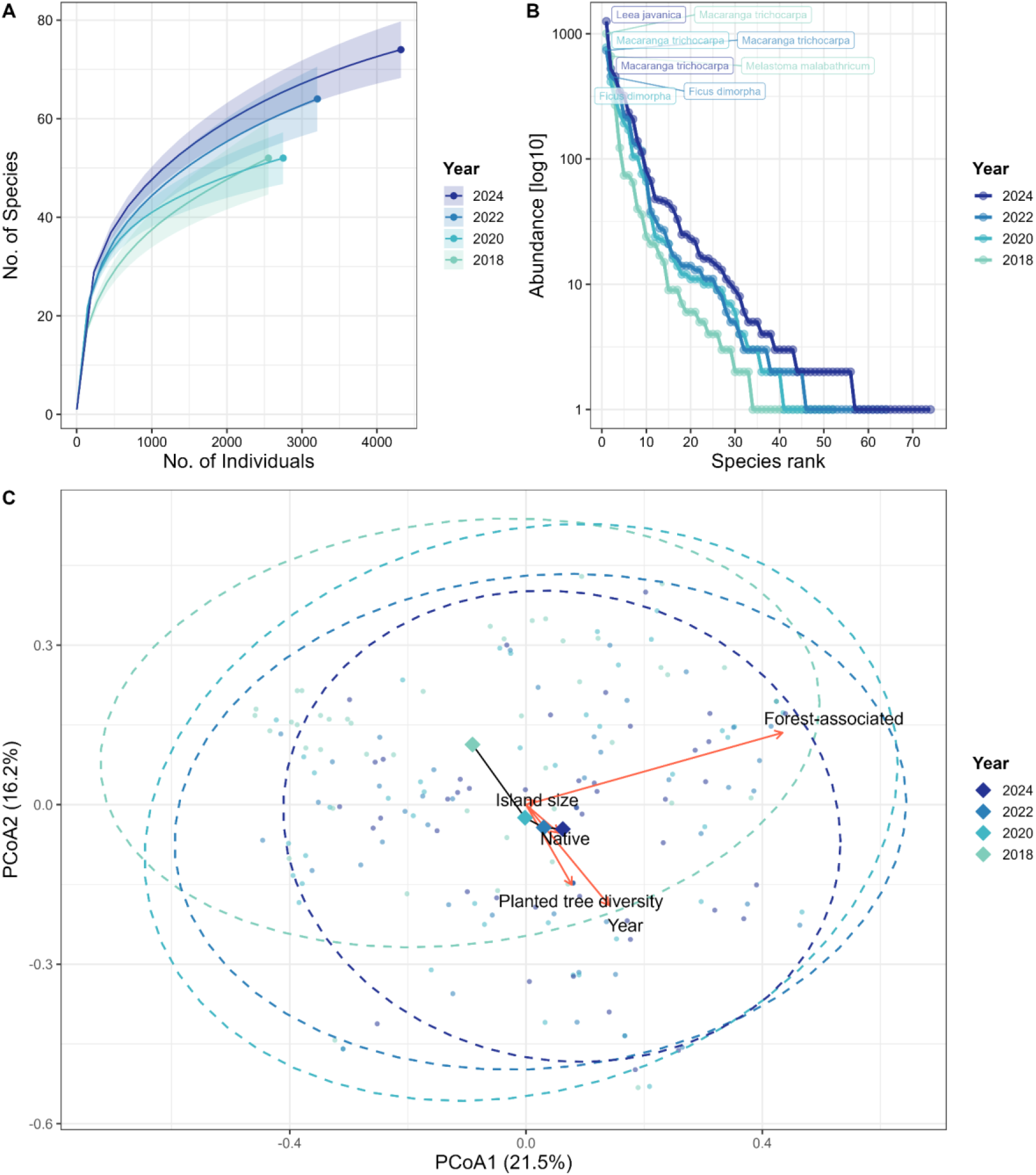
Temporal changes in the diversity and composition of naturally recruiting woody species within the EFForTS-BEE tree islands from 2018 to 2024. **(A)** Recruiting woody species taxonomic diversity at the landscape (gamma) scale based on sample-size rarefaction curves for 4 census years (2018, 2020, 2022, 2024) using ^0^*D*. Solid lines represent rarefaction curves based on the complete censuses, solid points represent observed recruiting diversity. Error bars represent 95% CIs from bootstrapping (*N* = 500 randomizations). **(B)** Rank-abundance curves for communities for 4 census years (2018, 2020, 2022, 2024), with the two most abundant species per year labelled. Abundances are displayed in log_10_-scale. **(C)** Principal coordinates analysis (PCoA) of recruiting community composition for 4 census years (2018, 2020, 2022, 2024). Points represent individual tree islands and dashed ellipses indicate 95% CIs around year groups. Large diamonds connected by a black line represent temporal shifts in year centroids. Red vectors indicate correlations between environmental variables and the ordination space; the proportion of forest-associated species, planted tree diversity, and year showed significant correlations with the PCoA axes.

Landscape-scale native richness increased strongly between 2020 and 2024 (^0^*D*; Fig. 2C), but slowly saturated for Shannon diversity (^1^*D*) or declined slightly for Simpson diversity (^2^*D*) (Fig. S3). In contrast, alien species diversity showed only a marginal increase for species richness (^0^*D*) and linearly decreased for Shannon diversity (^1^*D*) and Simpson diversity (^2^*D*) (Fig. 2C; Fig. S3), indicating that the overall rise in landscape-scale richness was driven primarily by native species. Alien abundance and alien basal area increased (Fig. S1), as a few species accounted for a disproportionately high share of individuals. In terms of basal area, recruits of oil palm (*Elaeis guineensis* Jacq., Arecaceae, alien) gained significant importance over the years, in 2024 accounting for 63 % of total basal area (Fig. S4B). The average proportion of native species per tree island remained high (83 - 88%).

Native forest-associated recruiting diversity increased steadily for all q orders (^0^*D,* ^1^*D,* ^2^*D*) at landscape scale, eventually exceeding native open-habitat recruiting diversity (Fig. 2D; Fig. S3). By 2024, 47% of all species were native forest-associated species, compared to 31% in 2018. Abundance remained higher for native open-habitat-associated species, while slowly saturating for native forest-associated species (Fig. S1). Basal area curves slowly saturated for both forest- and open-habitat-associated native species (Fig. S1). By 2024, native forest-associated species accounted for 20% of all individuals.

Community composition changed significantly through time (PERMANOVA: *F* = 4.32, *R*² = 0.069, *P* = 0.001), indicating temporal shifts in recruiting communities among tree islands. The PCoA showed that community centroids progressively shifted toward assemblages with greater representation of native and forest-associated species and individuals (Fig. 3C). Temporal changes were accompanied by increasing multivariate dispersion (*F* = 5.23, *P* = 0.002), indicating that islands became more diverse in their compositional trajectories through succession rather than converging toward a common community state. Overall, variation among tree islands was substantially greater than temporal variation, highlighting strong heterogeneity in community assembly dynamics across islands.

The ten most abundant species remained the same over time and accounted for ∼ 85% of individuals in 2024. Nevertheless, dominant species ranks shifted strongly between years (Fig. 3B; Fig. S4). In 2018, there were two dominant species at the landscape scale, *Macaranga trichocarpa* (Zoll.) Müll.Arg. (Euphorbiaceae, native) and *Melastoma malabathricum* L. (Melastomataceae, native), while in 2024, *Leea javanica* Blume (Vitaceae, native) was the only dominant species, which showed a steep increase in abundance (Fig. S4A).

Community evenness increased from 2018 to 2020, then slightly declined (Pielou’s evenness *J*_2018_ = 0.50, *J*_2020_ = 0.64, *J*_2022_ = 0.62, *J*_2024_ = 0.61). The number of singletons (i.e., species represented by a single individual) ranged between 12 and 19 and showed no directional trend over time. Communities were sampled at 99% sampling coverage across all years and q orders (Fig. S2B).

The increase in recruiting diversity at landscape (gamma) scale was also reflected at the local (alpha) scale, with the average species richness per tree island increasing over time (^0^*D*_2018_ = 6.12 ± 4.83 SD, ^0^*D*_2020_ = 6.50 ± 5.03 SD, ^0^*D*_2022_ = 7.60 ± 5.66 SD, ^0^*D*_2024_ = 9.13 ± 6.81 SD). Local evenness changed markedly over time, with some tree islands becoming more even as dominant species declined and more species were added, while other tree islands became more dominated by *Swietenia macrophylla* or *Leea javanica*. Yet, even in strongly dominated islands, there was a trend toward rising local species richness.

### 3.2 Effects of tree island area and planted tree diversity over time at the local scale

At the local (alpha) scale, year had a positive effect on recruiting diversity of native forest-associated species for ^0^*D* (*β*_2024_ = 0.46, *P* < .001, 95% confidence interval (CI) [0.37, 0.56]; Fig. 4; Table S4). Tree island area also increased local diversity of native forest-associated species, with effect sizes decreasing from ^0^*D* to ^2^*D* (*β* = 0.45 [^0^*D*], 0.36 [^1^*D*], 0.32 [^2^*D*]; Fig. S5; Table S4). No interaction between year and area was detected (Fig. 4; Fig. S5; Table S4). Planted tree diversity had a positive main effect on recruiting diversity of native forest-associated species for all q orders (*β*_0*D*_= 0.15, *P* = 0.015, CI [0.03, 0.26]; Fig. 4; Fig. S5; Table S4). However, there was a significant negative interaction between year and planted tree diversity, indicating declining influence of planted tree diversity through time for all q orders (*β*_2024,0*D*_ = -0.15, *P* = 0.002, CI [-0.24, -0.05]; Fig. 4; Fig. S5; Table S4). Overall, the mixed-effects models accounted for most of the variance (*R*^2^ _0*D*_= 86%, *R*^2^ _1*D*_= 81%, *R*^2^ _2*D*_= 76%), with most explained by fixed effects (*R*^2^_0*D*,fixed_ = 57%, *R*^2^_1*D*,fixed_ = 49%, *R*^2^_2*D*,fixed_ = 42%). The proportion of variance explained declined with increasing q orders. These results were also consistent when considering all native species (Fig. 4A; Fig. S5; Fig. S6; Table S5).

**Figure 4.**
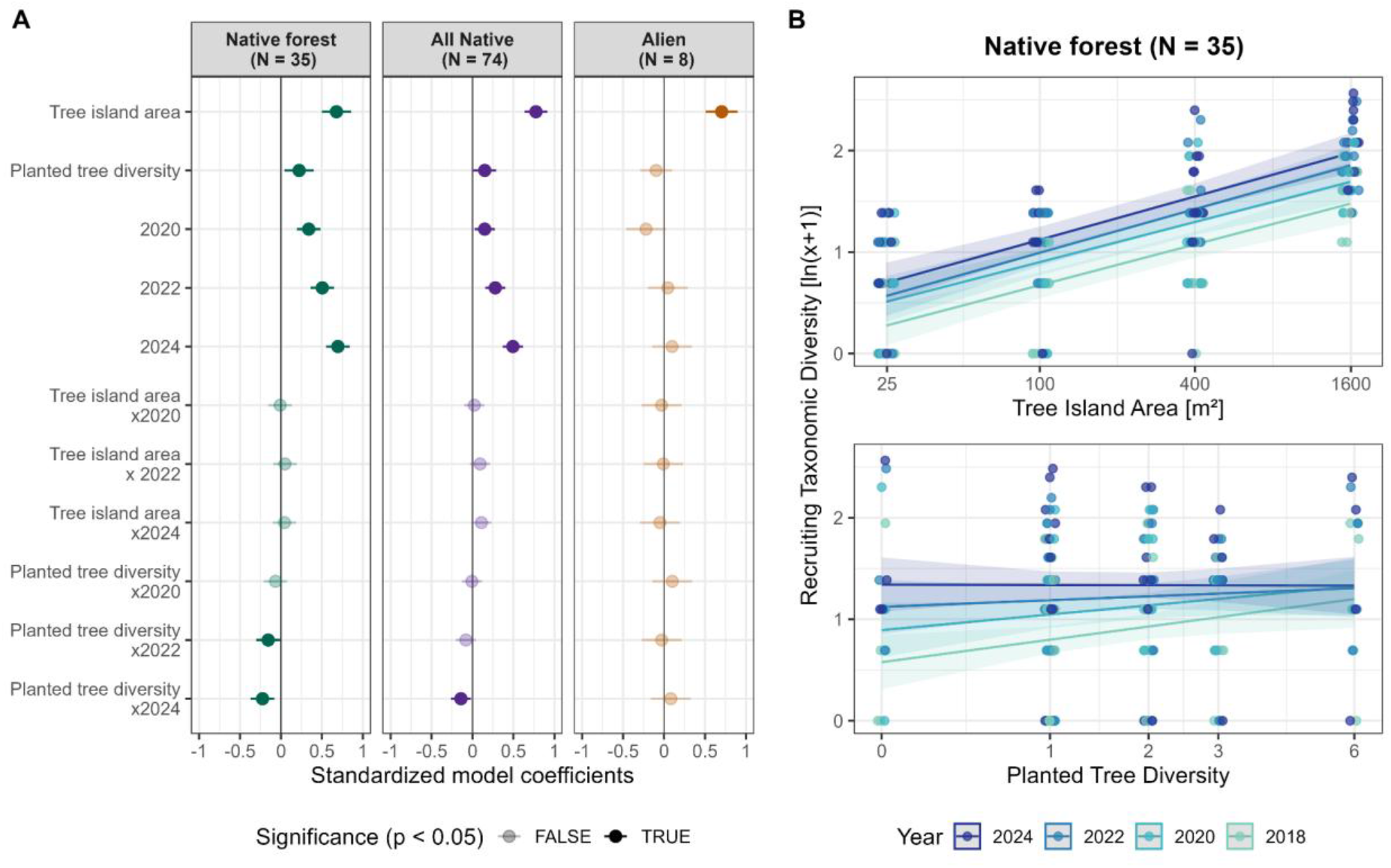
**(A)** Standardized coefficients from linear mixed-effects models testing the effects of tree island area, planted tree diversity, year, and their interactions on recruiting woody diversity (^0^*D*) for native forest-associated, native, and alien species. Filled circles indicate significant effects (*P* < 0.05), whereas transparent circles indicate non-significant effects (*P* > 0.05). Numbers in panel headers indicate total species richness within each group. **(B)** Model predictions showing the effects of tree island area (top) and planted tree diversity (bottom) on native forest-associated recruiting diversity (^0^*D*) for 4 census years (2018, 2020, 2022, 2024). Shaded areas represent 95% CIs. For model fitting, recruiting diversity was log-transformed as ln(x+1), while predictor variables were scaled and log-transformed as ln(x) for tree island area and ln(x+1) for planted tree diversity; x-axis labels show original predictor values.

In contrast to native recruiting species, the local diversity of alien recruiting species did not increase with time (Fig. 4A; Fig. S5; Fig. S7; Table S6). Tree island area was the only predictor that consistently increased local alien recruiting diversity across all q orders, while planted tree diversity showed a weak (non-significant) negative effect on alien recruiting diversity (Fig. 4A; Fig. S5; Fig. S7; Table S6).

## 4 Discussion

Forest restoration through natural regeneration is a long-term process (e.g., Brancalion et al., 2025; Gilman et al., 2016; Poorter et al., 2023), yet the extent to which native diversity can be recovered within industrial-scale plantation landscapes remains poorly understood, particularly over longer time frames (> 5 years) and under different restoration designs (Brancalion et al., 2025; Song et al., 2025). Our results demonstrate that tree islands embedded within industrial oil palm plantations promote sustained recovery of woody plant diversity. Recruiting species richness and abundance increased continuously, with no evidence of saturation. Importantly, the consistently high proportion of native species with progressive accumulation of forest-associated taxa, and the absence of alien-driven arrested succession, indicate that tree islands can facilitate the recovery of native plant diversity, even within heavily transformed agricultural landscapes.

At the local scale, restoration design strongly influenced recruitment trajectories of native and forest-associated species over time. Larger tree islands consistently supported higher native forest-associated recruiting diversity over time, most likely through a sampling effect, with additional contributions from area-dependent processes such as reduced edge effects and greater environmental heterogeneity. Because our censuses were complete, the sampling effect reflects the full local diversity rather than an artefact of incomplete sampling. In contrast, the lack of recruitment on multiple smaller tree islands in several census years suggests lower habitat stability of small tree islands, as small populations are more susceptible to stochastic processes and local extirpations, with a higher representation of transient species likely affecting also the ecosystem functions they provide (Allan et al., 2025). The positive effects of planted tree diversity were strongest during early stages and weakened over time, supporting the notion that natural recruitment progressively reduced initial differences in vegetation structure and microclimatic conditions among planting treatments. Whether these effects also extend to the functional and phylogenetic composition of recruiting communities remains to be tested.

### 4.1 Temporal changes in recruiting diversity and abundance

Recruiting diversity and abundance increased continuously over time at landscape scale, both across all species (^0^*D*) and among the common species (^1^*D*). These findings support our hypothesis (H1) that cumulative dispersal events and progressive vegetation development drive ongoing community assembly within tree islands (Poorter et al., 2023). Given the limited surrounding forest cover and the predominance of animal-dispersed species in the study system (Paterno et al., 2024), the continued increase in landscape-scale diversity suggests that long-distance seed dispersal remains a key recruitment mechanism (Piquer-Doblas et al., 2024; Poorter et al., 2023). The persistent occurrence of many singleton species further indicates ongoing species arrival, and non-saturation of recruiting diversity if arriving species are not transient but establishing stable coexisting populations over time (Alcántara et al., 2025; Allan et al., 2025).

Similar increases have been reported in tropical restoration experiments (Gilman et al., 2016; Zahawi et al., 2013), but recovery toward old-growth forests likely requires much longer periods and larger areas (Brancalion et al., 2025; Metz et al., 2026; Poorter et al., 2021): median recovery times in Neotropical secondary forests are estimated at five decades for species richness and centuries for species composition (Rozendaal et al., 2019).

The accumulation of genera and families over time further suggests increasing phylogenetic and functional diversity alongside taxonomic diversity, as ecological strategies are partly conserved within plant families (Cornwell et al., 2014), consistent with previous findings of correlated diversity facets in the EFForTS-BEE tree islands (Paterno et al., 2024) and broader evidence of increasing functional diversity during tropical forest succession (Van der Sande et al., 2024). Continued recruitment in tree islands may therefore contribute to ecosystem functioning and resilience (Ceulemans et al., 2019; De Bello et al., 2021; Garbowski et al., 2025; Wang et al., 2022).

At the local scale, recruiting diversity, abundance, and community composition varied strongly among tree islands, emphasizing stochasticity in community assembly and local restoration trajectories (cf. Brancalion et al., 2025; Norden et al., 2015; Zahawi et al., 2013). Such variation likely reflects differences in local site conditions, biotic interactions, life-history traits, and environmental stochasticity (Poorter et al., 2023), and can increase landscape-scale diversity through compositional differences among tree islands (Montoya-Sánchez et al., 2023).

### 4.2 Recovery of native forest-associated species

Supporting our second hypothesis (H2), native forest-associated diversity increased steadily over time at landscape scale, while the proportion of native open-habitat associated species declined, consistent with succession from open toward increasingly shaded conditions favoring forest-associated species (Poorter et al., 2023; Valladares et al., 2016).

In contrast, alien diversity showed no sustained increase over time, and declined with greater weight to common (^1^*D*) and highly abundant (^2^*D*) species, supporting our third hypothesis (H3) that alien recruiting diversity saturates earlier than native. This likely reflects the predominance of early-successional and limited representation of later-successional aliens in the regional species pool, with rapid colonization and fast growth traits facilitating their early establishment in productive environments (Gioria et al., 2023; Poorter et al., 2023). Although we found no evidence of alien-driven arrested succession (e.g., under local dominance of *Swietenia macrophylla*), persistent dominance of alien species could potentially cause arrested succession and declining native diversity (Matos et al., 2022), emphasizing the need for continued monitoring of future trajectories.

### 4.3 Long-term effects of experimental treatments

Tree island area consistently enhanced recruiting diversity, including native forest-associated species, consistent with the species-area relationship (Arrhenius, 1921) and equilibrium theory of island biogeography (MacArthur & Wilson, 1967; Rosenzweig, 2003), as well as previous evidence of greater plant recruitment in larger tree islands (Paterno et al., 2024; Zahawi & Augspurger, 2006; Zahawi et al., 2013). Contrary to our fourth hypothesis (H4), the area effect did not strengthen over time, indicating that available area strongly influenced recruitment from early succession onwards. In contrast, bird diversity in the EFForTS-BEE tree islands showed strengthening area effects over time (T. Geske, unpublished data), indicating an increasing importance of larger areas on recovering biodiversity. Nevertheless, small islands can still contribute to landscape diversity by hosting distinct species (Galán-Acedo et al., 2025; Montoya-Sánchez et al., 2023).

Higher planted tree diversity initially promoted recruiting diversity, particularly of native forest-associated species, but this effect weakened over time, supporting our fifth hypothesis (H5). Diverse plantings may initially have created more heterogeneous conditions that favoured a broader range of recruits, but as vegetation complexity subsequently increased through natural recruitment, conditions among tree islands gradually converged. Greater productivity, growth, and more stable survival in diverse tree plantings (Blondeel et al., 2024; Jensen et al., 2025; Zemp et al., 2019; Zheng et al., 2025), may have contributed to the initial positive effect of planted tree diversity by accelerating canopy closure, facilitating the establishment of more sensitive and shade-tolerant species (Gleiser et al., 2026). However, the temporal weakening of this effect may reflect a shift from strong early facilitation toward rising importance of niche competition as vegetation density increased (Poorter et al., 2023). Thus, unlike the strengthening tree diversity effects on productivity and other ecosystem functions reported elsewhere (e.g., Chen et al., 2025; Tatsumi, 2020; Zheng et al., 2024), tree diversity effects on natural regeneration did not strengthen over time in our study. Previous studies report mixed effects of planting treatments on natural recruitment (Gilman et al., 2016; Zahawi et al., 2013). Further, the weak negative effect of planted tree diversity on alien recruiting diversity observed in our study may suggest slightly greater resistance to alien establishment in more diverse plantings, consistent with biodiversity–invasibility theory (Elton, 1958; Halassy et al., 2023; Naeem et al., 2000; Perkins et al., 2011). Overall, restoration design was shown to shape successional trajectories with changing effects over time, underscoring the importance of long-term studies.

### 4.4 Implications for restoration planning and monitoring

Establishing tree islands with up to six native tree species in relatively small areas within the oil palm matrix, resulted in the natural recruitment of 82 woody plant species over 8 years, 35 of which native forest-associated. This finding provides compelling evidence that tree islands combined with natural regeneration can catalyze substantial biodiversity recovery within heavily modified plantation landscapes, highlighting their potential as a cost-efficient restoration strategy (Brancalion et al., 2025; Crouzeilles et al., 2020; Holl et al., 2020). Despite these gains, tree island diversity remained far below that of nearby lowland rainforests, which harbour >800 tree and shrub species, with very few alien taxa (Rembold et al., 2017). Recovering forests, particularly agroforestry systems, commonly remain less diverse than primary forests, frequently lacking later-successional, dispersal-limited, and rare species even after decades of regeneration (Brancalion et al., 2025; Metz et al., 2026; Song et al., 2025). Avoiding forest degradation in the first place therefore shows greater biodiversity benefits than restoration interventions alone can deliver (Miranda et al., 2026). Given the extensive area of oil palm plantations in Southeast Asia (Danylo et al., 2021), tree islands could substantially enhance biodiversity and connectivity across plantation landscapes, but remain a complementary strategy to, rather than a substitute for, conserving large, intact forest areas essential for sustaining native forest-associated diversity (Gibson et al., 2011; Norden et al., 2009; Paterno et al., 2024; Rembold et al., 2017; Watson et al., 2018).

As tree islands accumulate plant diversity and structural complexity (Kikuchi et al., 2024), they can provide habitat for multiple animal taxa and foster multi-taxa diversity at the landscape scale (Montoya-Sánchez et al., 2023; Zemp et al., 2023). Increasing bird diversity within the tree islands over the first decade, together with their attraction of birds to the wider landscape (T. Geske, unpublished data), and fruiting trees attracting long-tailed macaques (*Macaca fascicularis* Raffles, 1821), may further reinforce long-distance seed dispersal and recruitment of later-successional plant species (Albert et al., 2014; Corlett, 2017; Metz et al., 2026; Paterno et al., 2024; Poorter et al., 2023). These landscape-scale benefits could be further strengthened by actively reintroducing later-successional or dispersal-limited rare woody plant species (Cole et al., 2010; Holl et al., 2020; Holl et al., 2022; Joyce et al., 2025), controlling invasive species establishment (e.g., Halassy et al., 2023; Heleno et al., 2010; Matos et al., 2022; Weidlich et al., 2020), and allowing tree islands to expand into the surrounding oil palm through applied nucleation (e.g., Holl et al., 2020; Fig. S1 in Paterno et al., 2024; Werden et al., 2022; Zahawi & Augspurger, 2006), potentially providing a pathway for secondary forest regeneration, for example in protected-area buffer zones.

## 5 Conclusions

This study demonstrates that tree islands can promote sustained recovery of native forest-associated plant diversity in industrial oil palm landscapes over 8 years of natural regeneration, without evidence of alien-driven arrested succession. Initial restoration design shaped recruitment trajectories over time: larger tree islands consistently supported greater diversity of native forest-associated species, whereas the positive effects of planted tree diversity were strongest during earlier stages of succession. Despite substantial biodiversity gains, tree islands recovered only a fraction of the plant diversity found in nearby lowland rainforests. Tree islands therefore offer a promising, scalable restoration strategy for enhancing biodiversity in oil palm landscapes, while complementing - not replacing - the strict protection of remaining tropical forests.

## Authors’ contributions

Vivien Michel, Gustavo Brant Paterno, Fabian Brambach, and Holger Kreft conceived the study; Vivien Michel led the field data collection and data curation, with contributions from Fabian Brambach and Gustavo Brant Paterno; Vivien Michel and Gustavo Brant Paterno developed the analytical framework and Vivien Michel performed the data analysis and produced the figures; Vivien Michel wrote the first draft of the manuscript; Holger Kreft, Dirk Hölscher, Bambang Irawan, and Leti Sundawati contributed to project administration and funding acquisition; Gustavo Brant Paterno and Holger Kreft supervised the work. All authors contributed critically to manuscript revisions and approved the final version for publication.

## Statement on inclusion

Our study builds on more than a decade of Indonesian–German collaboration and includes scientists based in Indonesia, where the research was conducted. Local collaborators were involved in study implementation, fieldwork, and interpretation of the results.

## Acknowledgements

We sincerely appreciate the Ministry of Research and Technology of Indonesia (RISTEK), the Indonesian Institute of Sciences (LIPI), the Ministry of Forestry (PHKA) and Restoration Ecosystem Indonesia (PT REKI) for granting research permits (FB 232/SIP/IV/FR/10/2022; GBP 100/SIP/IV/FR/2/2023; VM 551/SIP/IV/FR/7/2024). We also extend our gratitude to our colleagues and field assistants for their invaluable support during fieldwork and laboratory analyses. We thank PT Humusindo for granting access to and use of their properties. We thank Juliandi, K. H. Dalimunthe, E. J. Siahaan, E. Mauliarta and T. Simanjuntak for their support in field campaigns. We thank Clara Zemp for providing data on natural regeneration from the first two censuses. The EFForTS-BEE experiment is part of TreeDivNet, a global network of tree diversity experiments (https://treedivnet.ugent.be/).

## Funding information

The establishment and maintenance of EFForTS-BEE was funded by the Deutsche Forschungsgemeinschaft (DFG), project number 192626868–SFB 990 in the framework of the collaborative German-Indonesian research project CRC990/EFForTS as well as the follow-on bundle of projects “Conversion of rainforest into rubber and oil palm plantations in Sumatra, Indonesia: Landscape-scale changes in biodiversity and ecosystem functioning, and restoration pathways”, project number 532776526 grants HO 2119/8-1 and KR 3974/13-1. N.G.R and H.K. further acknowledge funding by Biodiversa+, the European Biodiversity Partnership, in the context of the BioMonI – Biodiversity monitoring of island ecosystems project under the 2022-2023 BiodivMon joint call. It was co-funded by the European Commission (GA No. 101052342) and the Deutsche Forschungsgemeinschaft (DFG, German Research Foundation)—project ID 533271599 for the University of Göttingen (Germany).

## Conflict of interest

The authors declare no conflicts of interest.

## Data availability statement

All data and code necessary to reproduce the analyses and figures presented in this study will be made available in the GRO.data repository.

## Notes

### Competing Interest Statement

The authors have declared no competing interest.

