## Supplementary materials for "Natural regeneration in tree islands drives long-term recovery of native forest-associated woody species in oil palm landscapes"

### **Supplementary figures**

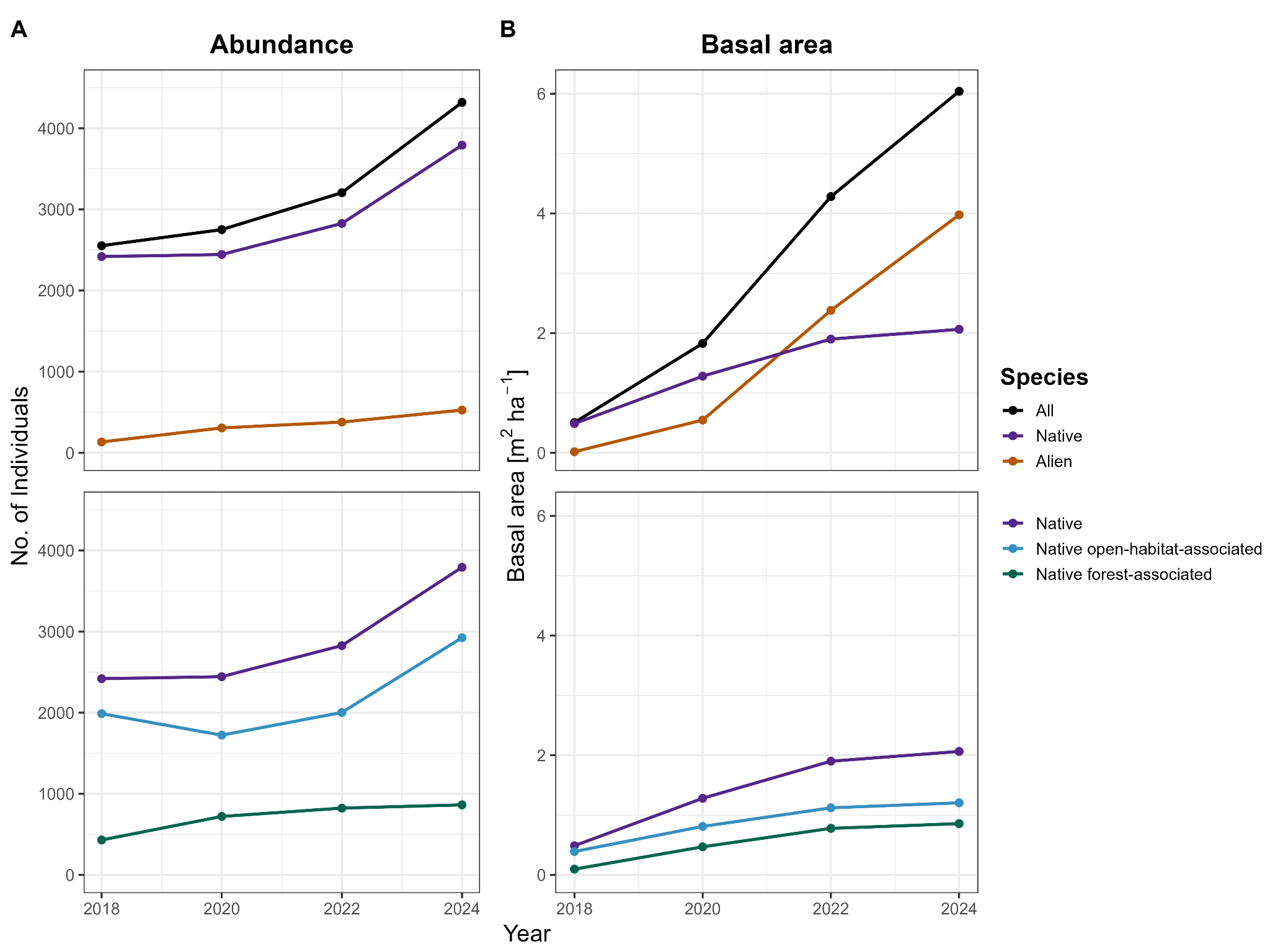

**Figure S1:** Recruiting abundance **(A)** and basal area **(B)** of native versus alien species (top) and habitat types (bottom) of native species (open habitats, forest) over 4 census years (2018, 2020, 2022, 2024).

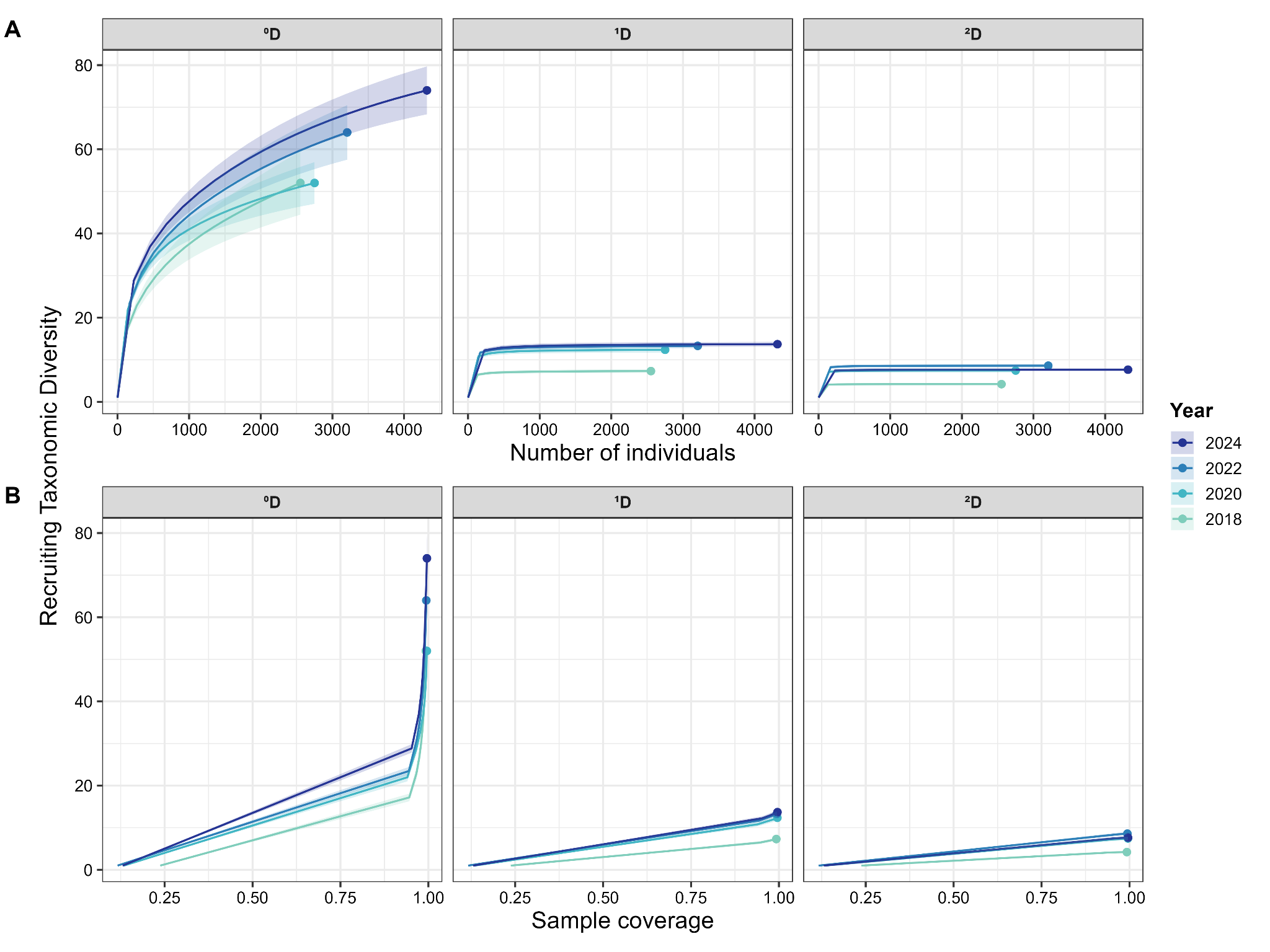

**Figure S2**: Recruiting taxonomic diversity according to **(A)** sample-size based rarefaction and **(B)** coverage-based rarefaction for 4 census years (2018, 2020, 2022, 2024) and three q orders (^0^*D,* ^1^*D,* ^2^*D*) at landscape scale. Solid lines represent rarefaction curves based on the complete censuses; solid points represent observed recruiting diversity. Error bars represent 95% CIs from bootstrapping (*N* = 500 randomizations).

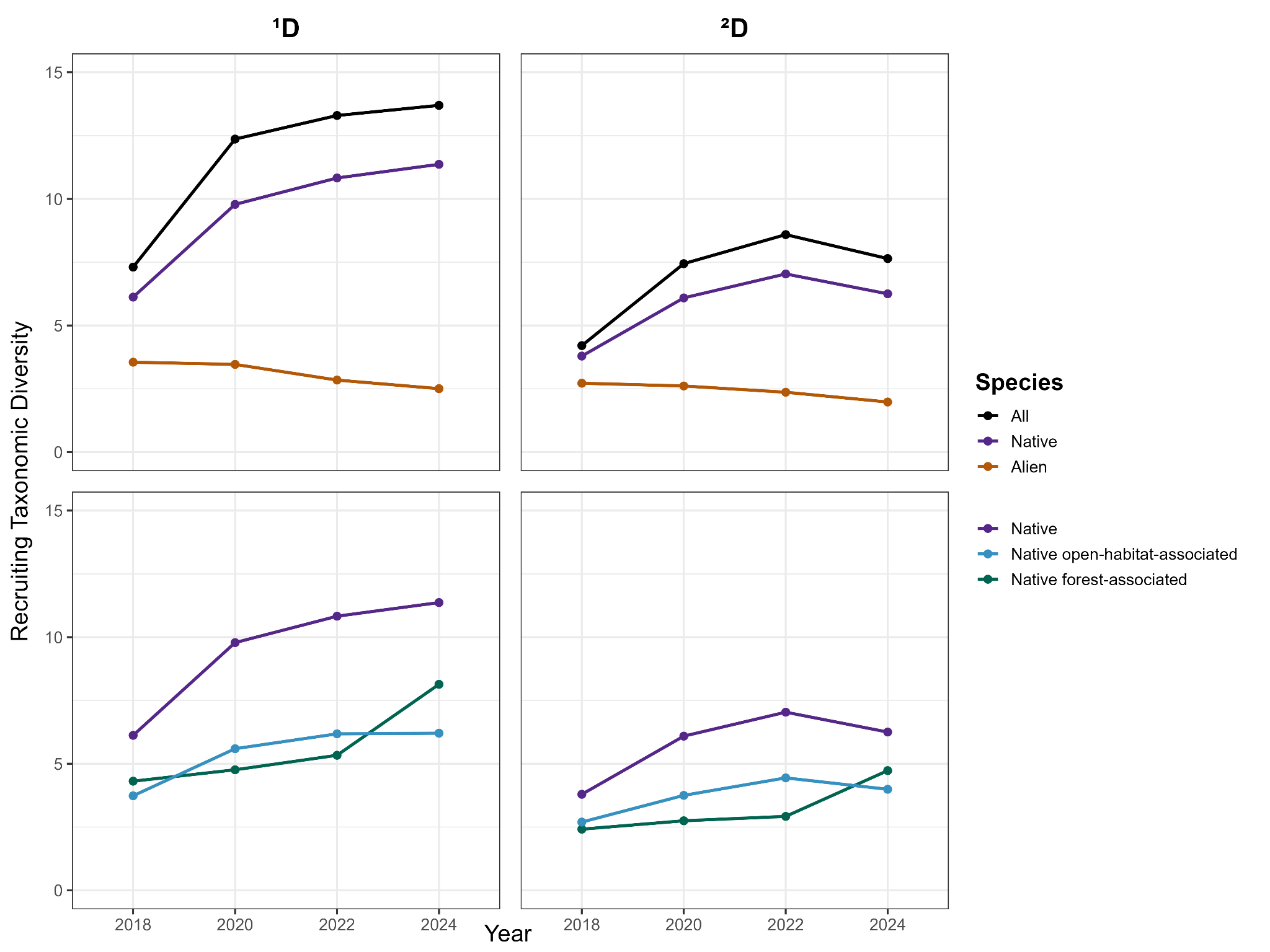

**Figure S3:** Recruiting taxonomic diversity of native versus alien species (top) and habitat types (bottom) of native species (open habitats, forest) over 4 census years (2018, 2020, 2022, 2024) for ^1^*D* and ^2^*D*.

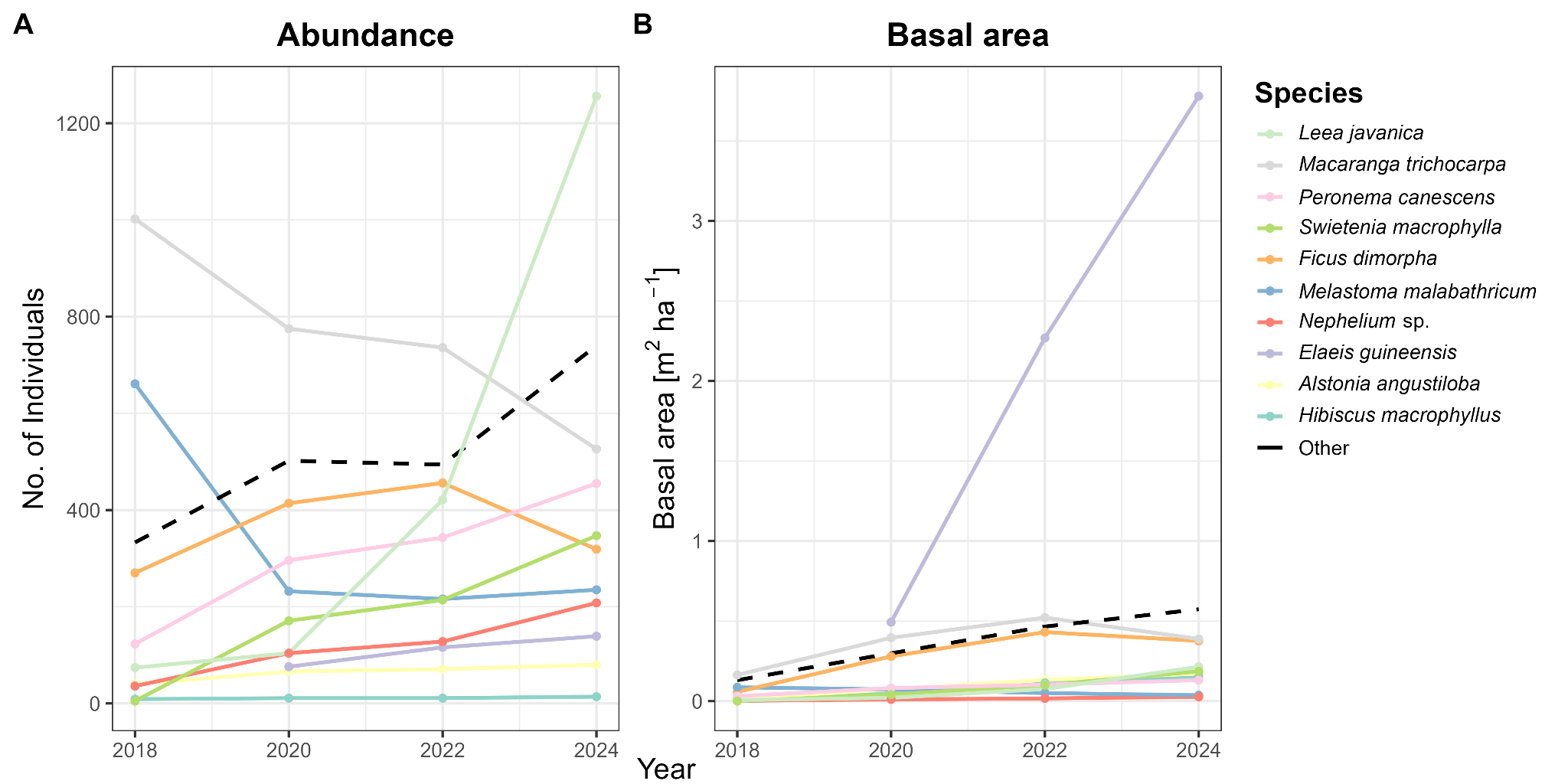

**Figure S4:** Trajectories of abundance **(A)** and basal area **(B)** over 4 census years (2018, 2020, 2022, 2024) for the ten species ranking highest in abundance or basal area in 2024. The dashed line represents the combined total of all remaining species. The legend displays species in decreasing order based on 2024 abundance.

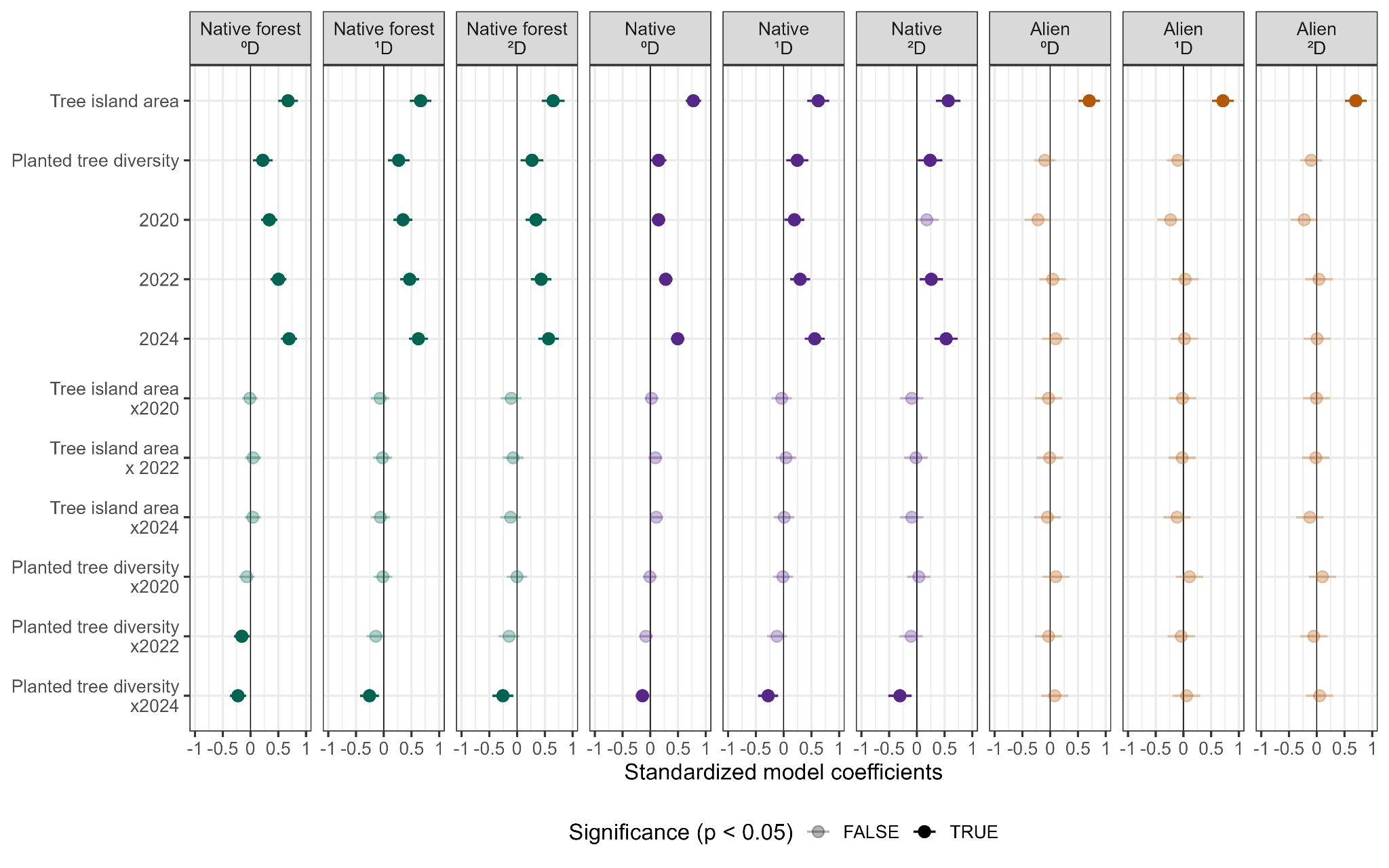

**Figure S5:** Standardized model coefficients for native forest-associated recruits (left), native recruits (center), and alien recruits (right) at three q orders (^0^*D,* ^1^*D,* ^2^*D*). Transparent circles represent non-significant predictors (*P* > 0.05).

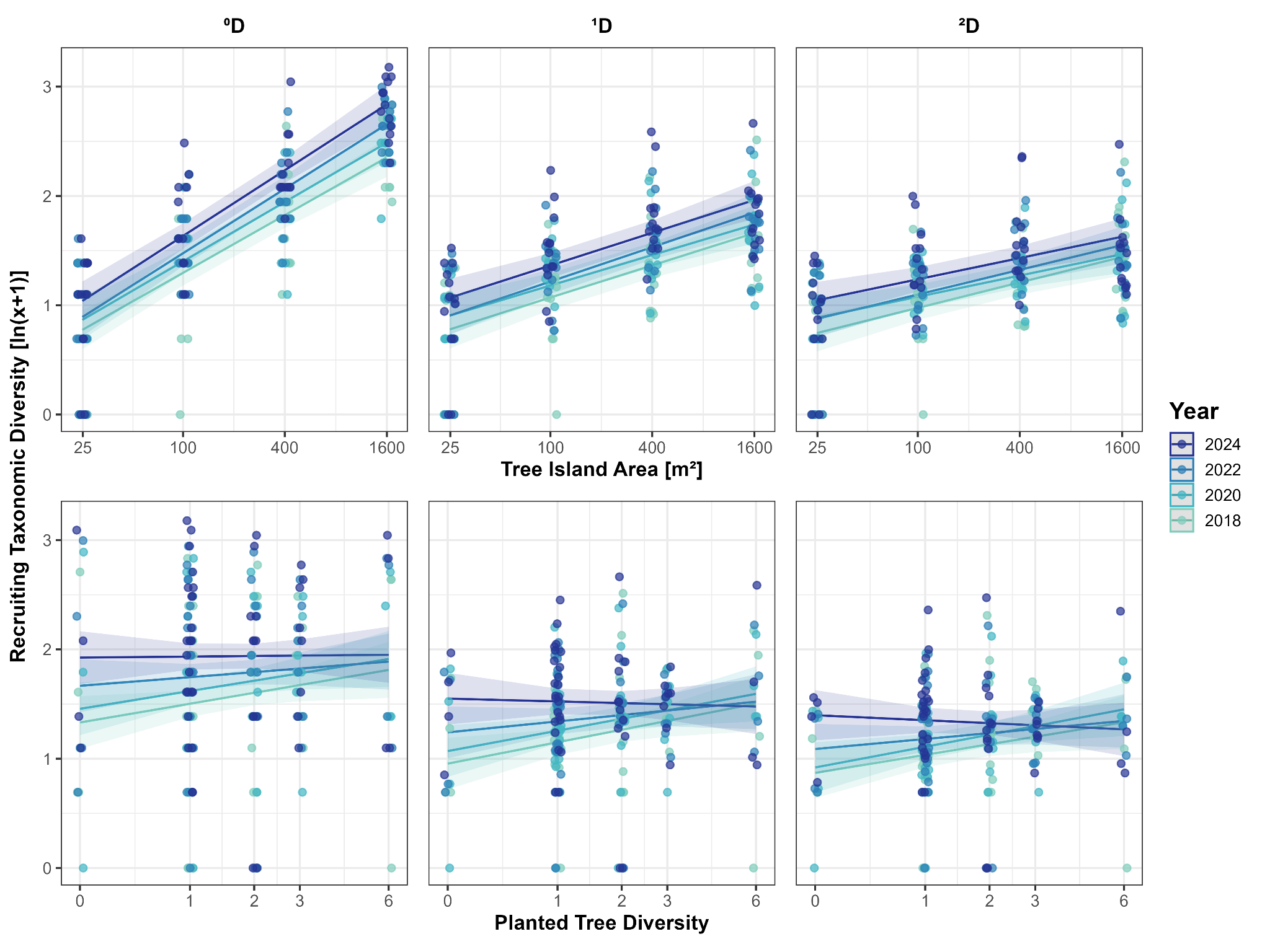

**Figure S6:** LMM predictions for tree island area (top) and planted tree diversity (bottom) effects on native recruiting diversity (^0^*D* = 74) at three q orders (^0^*D,* ^1^*D,* ^2^*D*) over 4 census years (2018, 2020, 2022, 2024). For model fitting, recruiting diversity was log-transformed as ln(x+1), while predictor variables were scaled and log-transformed as ln(x) for tree island area and ln(x+1) for planted tree diversity; x-axis labels show original predictor values.

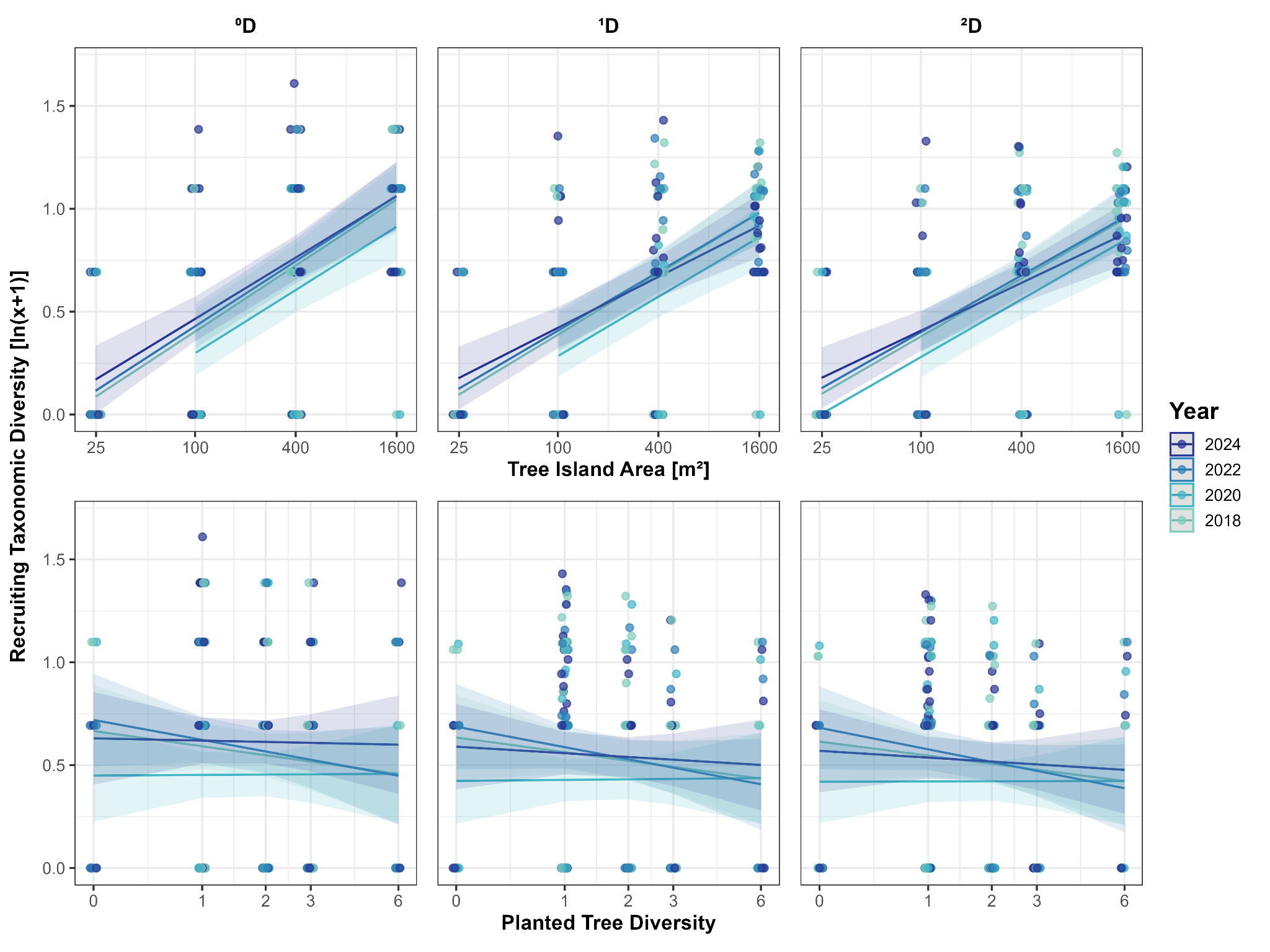

**Figure S7**: LMM predictions for tree island area (top) and planted tree diversity (bottom) effects on alien recruiting diversity (^0^*D* = 8) at three q orders (^0^*D,* ^1^*D,* ^2^*D*) for 4 census years (2018, 2020, 2022, 2024). For model fitting, recruiting diversity was log-transformed as ln(x+1), while predictor variables were scaled and log-transformed as ln(x) for tree island area and ln(x+1) for planted tree diversity; x-axis labels show original predictor values.

### **Supplementary tables**

**Table S1**: Mechanistic hypotheses (H1- H5) for the three subsets of all, alien, and native forest-associated species, as well as the experimental treatments of tree island area and planted tree diversity.

|  | **Species subset/**  **Experimental treatment** | **Hypothesis** | **Mechanism** |
| --- | --- | --- | --- |
| ***H1*** | ***All species*** | General increase over time   a) Steady increase b) Saturation c) Slight decline | Cumulative seed arrival,  Vegetation development  Long-distance dispersal by animals  Limited regional species pool  Competitive exclusion,  Suppression by dominant alien species |
| ***H2*** | ***Alien species*** | Early increase and saturation / decline | Rapid colonization,  Smaller regional species pool |
| ***H3*** | ***Native forest-associated species*** | Delayed increase and saturation / decline | More favorable conditions at later-successional stages |
| ***H4*** | ***Larger tree island area*** | Higher recruiting diversity with strengthening effect over time | Species-area relationship,  Reduced edge effects,  Greater habitat heterogeneity |
| ***H5*** | ***Higher planted tree diversity*** | Higher recruiting diversity initially with diminishing effect over time | Greater habitat heterogeneity,  Converging light and microclimatic conditions over time through natural recruitment |

**Table S2**: List of recruiting liana species in 2024, with respective lifeform, origin, habitat, 2024 abundance, and tree island occupancy (i.e., percentage of tree islands in which the species is present). Families and respective species are listed in alphabetical order.

| Family | Species | Lifeform | Origin | Habitat | 2024 | Occupancy  2024 (%) |
| --- | --- | --- | --- | --- | --- | --- |
| Annonaceae | *Uvaria* cf. *grandiflora* Roxb. ex Hornem. | liana | native | forest | 1 | 1.92 |
| Apocynaceae | *Chonemorpha verrucosa* (Blume) D.J.Middleton | liana | native | forest | 22 | 5.77 |
| Apocynaceae | *Secamone* sp.KR5266 | liana | native | NA | 50 | 21.15 |
| Celastraceae | *Reissantia cassinoides* (DC.) Ding Hou | liana | native | forest | 7 | 9.62 |
| Combretaceae | *Combretum* cf. *sundaicum* Miq. | liana | native | open | 63 | 25.00 |
| Convolvulaceae | *Bonamia semidigyna* (Roxb.) Hallier f. | liana | native | open | 9 | 3.85 |
| Convolvulaceae | *Camonea pilosa* (Houtt.) A.R.Simões & Staples | liana | native | open | 90 | 15.38 |
| Convolvulaceae | *Erycibe rheedei* Blume | liana | native | open | 2 | 3.85 |
| Dilleniaceae | *Tetracera scandens* (L.) Merr. | liana | native | open | 2 | 1.92 |
| Fabaceae | *Calopogonium mucunoides* Desv. | liana | alien | open | 3 | 1.92 |
| Fabaceae | *Centrosema molle* Mart. ex Benth. | liana | alien | open | 8 | 11.54 |
| Fabaceae | *Derris amoena* Benth. | liana | native | open | 7 | 1.92 |
| Fabaceae | *Mezoneuron sumatranum* (Roxb.) Wight & Arn. ex Voigt | liana | native | open | 1 | 1.92 |
| Fabaceae | *Millettia sericea* (Vent.) Wight & Arn. ex Hassk. | liana | native | open | 2 | 1.92 |
| Fabaceae | *Mucuna biplicata* Teijsm. & Binn. ex Kurz | liana | native | open | 26 | 21.15 |
| Fabaceae | *Phanera semibifida* (Roxb.) Benth. | liana | native | open | 36 | 25.00 |
| Gnetaceae | *Gnetum tenuifolium* Ridl. | liana | native | forest | 2 | 3.85 |
| Melastomataceae | *Macrolenes* cf. *dimorpha* (Craib) J.F.Maxwell | liana | native | open | 3 | 1.92 |
| Menispermaceae | *Arcangelisia flava* (L.) Merr. | liana | native | forest | 1 | 1.92 |
| Oleaceae | *Jasminum elongatum* (P.J.Bergius) Willd. | liana | native | open | 1 | 1.92 |
| Phyllanthaceae | *Bridelia cinnamomea* Hook.f. | liana | native | forest | 3 | 1.92 |
| Primulaceae | *Maesa sumatrana* Scheff. | liana | native | open | 4 | 3.85 |
| Resedaceae | *Stixis scortechinii* (King) M.Jacobs | liana | native | open | 5 | 5.77 |
| Rubiaceae | *Coptosapelta* cf. *tomentosa* (Blume) Valeton ex K.Heyne | liana | native | forest | 6 | 1.92 |
| Rubiaceae | *Mussaenda frondosa* L. | liana | native | open | 4 | 5.77 |
| Rubiaceae | *Uncaria attenuata* Korth. | liana | native | open | 24 | 21.15 |
| Rubiaceae | *Uncaria cordata* (Lour.) Merr. | liana | native | open | 17 | 23.08 |
| Vitaceae | *Tetrastigma coriaceum* (DC.) Gagnep. | liana | native | open | 9 | 5.77 |

**Table S3**: List of all recruiting woody species recorded between 2018 and 2024, with respective lifeform, origin, habitat, 2018-2024 occurrences, and tree island occupancy in 2024 (i.e., percentage of tree islands in which the species was present). Families and their respective species are listed in alphabetical order.

| Family | Species | Origin | Lifeform | Habitat | 2018 | 2020 | 2022 | 2024 | Occupancy  2024 (%) |
| --- | --- | --- | --- | --- | --- | --- | --- | --- | --- |
| Annonaceae | *Polyalthia cauliflora* Hook.f. & Thomson | native | tree | forest | 0 | 0 | 1 | 1 | 1.92 |
| Apocynaceae | *Alstonia angustiloba* Miq. | native | tree | forest | 40 | 66 | 71 | 80 | 50 |
| Apocynaceae | *Tabernaemontana pauciflora* Blume | native | treelet | forest | 6 | 22 | 21 | 33 | 23.08 |
| Arecaceae | *Elaeis guineensis* Jacq. | alien | tree | open | 0 | 76 | 116 | 139 | 46.15 |
| Asteraceae | *Clibadium surinamense* L. | alien | shrub | open | 21 | 6 | 1 | 0 | 0 |
| Cannabaceae | *Trema tomentosum* (Roxb.) H.Hara | native | tree | open | 65 | 12 | 1 | 0 | 0 |
| Combretaceae | *Terminalia* cf. *calamansanai* (Blanco) Rolfe | native | tree | open | 1 | 1 | 1 | 1 | 1.92 |
| Dilleniaceae | *Dillenia* cf. *excelsa* (Jack) Martelli ex Gilg. | native | tree | forest | 6 | 11 | 11 | 16 | 3.85 |
| Dipterocarpaceae | *Hopea sangal* Korth. | native | tree | forest | 0 | 0 | 1 | 1 | 1.92 |
| Elaeocarpaceae | *Elaeocarpus cf. stipularis* Blume | native | tree | forest | 0 | 0 | 1 | 2 | 3.85 |
| Euphorbiaceae | *Croton argyratus* Blume | native | tree | forest | 6 | 10 | 16 | 25 | 19.23 |
| Euphorbiaceae | *Macaranga* aff. *conifera* (Rchb.f. & Zoll.) Müll.Arg. | native | tree | forest | 15 | 10 | 14 | 15 | 15.38 |
| Euphorbiaceae | *Macaranga bancana* (Miq.) Müll.Arg. | native | tree | open | 1 | 1 | 6 | 17 | 15.38 |
| Euphorbiaceae | *Macaranga gigantea* (Rchb.f. & Zoll.) Müll.Arg. | native | tree | open | 2 | 1 | 1 | 2 | 3.85 |
| Euphorbiaceae | *Macaranga hosei* King ex Hook.f. | native | tree | open | 1 | 0 | 0 | 0 | 0 |
| Euphorbiaceae | *Macaranga trichocarpa* (Zoll.) Müll.Arg. | native | shrub | open | 1002 | 775 | 736 | 526 | 48.08 |
| Euphorbiaceae | *Mallotus macrostachyus* (Miq.) Müll.Arg. | native | tree | open | 3 | 4 | 3 | 4 | 3.85 |
| Euphorbiaceae | *Mallotus paniculatus* (Lam.) Müll.Arg. | native | tree | open | 2 | 2 | 2 | 0 | 0 |
| Euphorbiaceae | *Mallotus peltatus* (Geiseler) Müll.Arg. | native | tree | open | 7 | 10 | 27 | 67 | 21.15 |
| Euphorbiaceae | *Triadica cochinchinensis* Lour. | native | tree | open | 0 | 0 | 0 | 1 | 1.92 |
| Fabaceae | *Archidendron jiringa* (Jack) I.C.Nielsen | native | tree | forest | 1 | 1 | 1 | 11 | 5.77 |
| Fabaceae | *Dalbergia junghuhnii* Benth. | native | treelet | open | 0 | 0 | 0 | 1 | 1.92 |
| Fabaceae | *Peltophorum dasyrhachis* (Miq.) Kurz | native | tree | forest | 0 | 0 | 0 | 1 | 1.92 |
| Fabaceae | *Senna alata* (L.) Roxb. | alien | shrub | open | 9 | 14 | 5 | 10 | 1.92 |
| Hypericaceae | *Cratoxylum formosum* (Jack) Benth. & Hook.f. ex Dyer | native | tree | forest | 2 | 3 | 2 | 2 | 3.85 |
| Hypericaceae | *Cratoxylum sumatranum* (Jack) Blume | native | tree | forest | 0 | 0 | 3 | 3 | 3.85 |
| Lamiaceae | *Callicarpa pentandra* Roxb. | native | tree | open | 1 | 0 | 0 | 0 | 0 |
| Lamiaceae | *Clerodendrum disparifolium* Blume | native | tree | forest | 17 | 21 | 28 | 46 | 21.15 |
| Lamiaceae | *Peronema canescens* Jack. | native | tree | open | 123 | 296 | 343 | 455 | 34.62 |
| Lamiaceae | *Vitex quinata* (Lour.) F.N.Williams | native | tree | forest | 21 | 36 | 33 | 40 | 21.15 |
| Lauraceae | *Litsea* cf. *robusta* Blume | native | tree | forest | 0 | 0 | 2 | 3 | 3.85 |
| Lauraceae | *Litsea umbellata* (Lour.) Merr. | native | tree | open | 1 | 3 | 14 | 48 | 34.62 |
| Malvaceae | *Commersonia bartramia* (L.) Merr. | native | tree | open | 1 | 1 | 1 | 1 | 1.92 |
| Malvaceae | *Hibiscus macrophyllus* Roxb. ex Hornem. | native | tree | open | 9 | 11 | 11 | 14 | 13.46 |
| Malvaceae | *Trichospermum javanicum* Blume | native | tree | forest | 1 | 0 | 0 | 0 | 0 |
| Malvaceae | *Urena lobata* L. | native | shrub | open | 1 | 0 | 1 | 2 | 1.92 |
| Melastomataceae | *Bellucia pentamera* Naudin | alien | tree | open | 1 | 16 | 5 | 6 | 1.92 |
| Melastomataceae | *Melastoma malabathricum* L. | native | shrub | open | 661 | 232 | 216 | 235 | 40.38 |
| Melastomataceae | *Pternandra azurea* (Blume) Burkill | native | tree | open | 0 | 0 | 1 | 2 | 1.92 |
| Meliaceae | *Swietenia macrophylla* King | alien | tree | forest | 5 | 171 | 214 | 347 | 23.08 |
| Moraceae | *Artocarpus dadah* Miq. | native | tree | forest | 0 | 0 | 1 | 1 | 1.92 |
| Moraceae | *Artocarpus heterophyllus* Lam. | alien | tree | forest | 0 | 0 | 0 | 1 | 1.92 |
| Moraceae | *Artocarpus limpato* Miq. | native | tree | forest | 0 | 0 | 0 | 1 | 1.92 |
| Moraceae | *Ficus aurata* (Miq.) Miq. | native | treelet | open | 4 | 4 | 3 | 16 | 5.77 |
| Moraceae | *Ficus caulocarpa* (Miq.) Miq. | native | tree | forest | 1 | 2 | 2 | 4 | 7.69 |
| Moraceae | *Ficus* cf. *padana* Burm.f. | native | tree | open | 0 | 1 | 1 | 0 | 0 |
| Moraceae | *Ficus dimorpha* King | native | treelet | forest | 270 | 414 | 456 | 319 | 69.23 |
| Moraceae | *Ficus glandulifera* (Wall. ex Miq.) King | native | tree | forest | 4 | 2 | 3 | 2 | 3.85 |
| Moraceae | *Ficus grossularioides* Burm.f. | native | tree | open | 1 | 0 | 0 | 0 | 0 |
| Moraceae | *Ficus variegata* Blume | native | tree | open | 5 | 11 | 11 | 5 | 7.69 |
| Moraceae | *Ficus vrieseana* Miq. | native | tree | open | 1 | 9 | 13 | 25 | 13.46 |
| Moraceae | *Sloetia elongata* (Miq.) Koord. | native | tree | open | 1 | 1 | 3 | 3 | 5.77 |
| Myrtaceae | *Syzygium* sp.VM0002 | native | tree | NA | 0 | 0 | 0 | 3 | 3.85 |
| Opiliaceae | *Champereia manillana* (Blume) Merr. | native | treelet | forest | 0 | 2 | 2 | 2 | 3.85 |
| Pandaceae | *Galearia filiformis* (Blume) Boerl. | native | treelet | forest | 0 | 1 | 2 | 5 | 5.77 |
| Pandaceae | *Microdesmis caseariifolia* Planch. | native | treelet | forest | 3 | 9 | 13 | 13 | 17.31 |
| Pentaphylacaceae | *Eurya nitida* Korth. | native | treelet | open | 1 | 1 | 1 | 1 | 1.92 |
| Phyllanthaceae | *Aporosa benthamiana* Hook.f. | native | tree | forest | 0 | 0 | 1 | 1 | 1.92 |
| Phyllanthaceae | *Aporosa octandra* (Buch.-Ham. ex D.Don) Vickery | native | tree | open | 0 | 0 | 0 | 1 | 1.92 |
| Phyllanthaceae | *Baccaurea brevipes* Hook.f. | native | treelet | forest | 0 | 0 | 1 | 2 | 3.85 |
| Phyllanthaceae | *Breynia racemosa* (Blume) Müll.Arg. | native | treelet | open | 9 | 23 | 14 | 47 | 25 |
| Phyllanthaceae | *Bridelia glauca* Blume | native | tree | forest | 0 | 1 | 1 | 2 | 3.85 |
| Phyllanthaceae | *Glochidion borneense* (Müll.Arg.) Boerl. | native | tree | open | 1 | 0 | 1 | 2 | 3.85 |
| Poaceae | *Gigantochloa scortechinii* Gamble | native | shrub | open | 0 | 193 | 134 | 113 | 26.92 |
| Rhizophoraceae | *Carallia brachiata* (Lour.) Merr. | native | tree | forest | 0 | 0 | 0 | 1 | 1.92 |
| Rhizophoraceae | *Gynotroches axillaris* Blume | native | tree | open | 0 | 0 | 2 | 2 | 1.92 |
| Rubiaceae | *Adina eurhyncha* (Miq.) Å.Krüger & Löfstrand | native | tree | forest | 0 | 0 | 0 | 2 | 3.85 |
| Rubiaceae | *Canthium horridum* Baill. | native | shrub | open | 3 | 7 | 8 | 8 | 7.69 |
| Rubiaceae | *Eumachia cf. montana* (Blume) I.M.Turner | native | shrub | forest | 0 | 1 | 4 | 9 | 11.54 |
| Rubiaceae | *Ixora* sp.VM0025 | native | shrub | NA | 0 | 0 | 0 | 1 | 1.92 |
| Rubiaceae | *Neolamarckia cadamba* (Roxb.) Bosser | native | tree | open | 1 | 1 | 1 | 1 | 1.92 |
| Rubiaceae | *Neonauclea calycina* (Bartl. ex DC.) Merr. | native | tree | open | 4 | 7 | 9 | 23 | 13.46 |
| Rubiaceae | *Psychotria viridiflora* Reinw. ex Blume | native | shrub | forest | 0 | 0 | 0 | 5 | 7.69 |
| Rubiaceae | *Urophyllum peltistigma* Miq. | native | treelet | forest | 0 | 3 | 3 | 4 | 5.77 |
| Rutaceae | *Clausena excavata* Burm.f. | native | tree | open | 1 | 12 | 17 | 44 | 30.77 |
| Salicaceae | *Homalium caryophyllaceum* (Zoll. & Moritzi) Benth. | native | treelet | forest | 2 | 2 | 2 | 3 | 1.92 |
| Sapindaceae | *Nephelium* sp.DMF0033 | native | tree | forest | 36 | 104 | 128 | 208 | 30.77 |
| Simaroubaceae | *Brucea javanica* (L.) Merr. | native | treelet | open | 1 | 0 | 0 | 1 | 1.92 |
| Solanaceae | *Solanum jamaicense* Mill. | alien | shrub | open | 74 | 24 | 38 | 22 | 32.69 |
| Strombosiaceae | *Strombosia ceylanica* Gardner | native | tree | forest | 0 | 0 | 0 | 1 | 1.92 |
| Verbenaceae | *Stachytarpheta australis* Moldenke | alien | shrub | open | 24 | 0 | 0 | 2 | 3.85 |
| Vitaceae | *Leea javanica* Blume | native | tree | open | 74 | 104 | 421 | 1256 | 51.92 |

**Table S4**: LMM outputs for native forest-associated recruiting diversity at three q orders (^0^*D,* ^1^*D,* ^2^*D*).

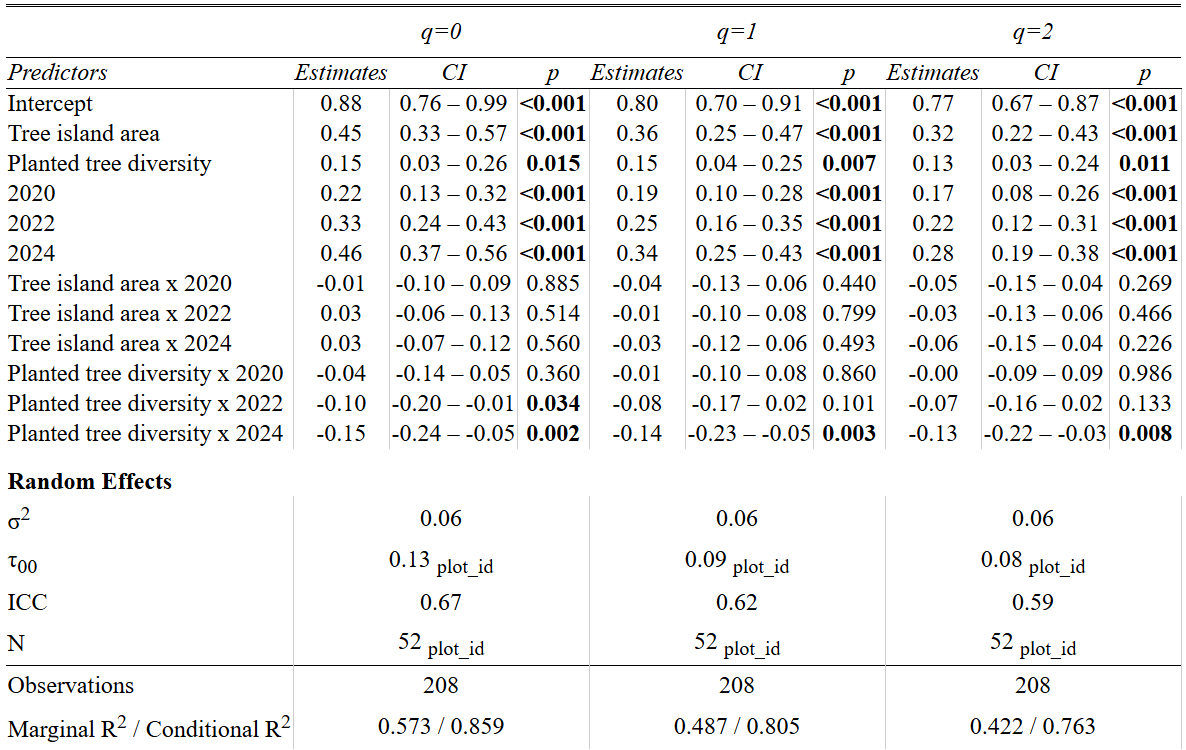

**Table S5**: LMM outputs for native recruiting diversity at three q orders (^0^*D,* ^1^*D,* ^2^*D*).

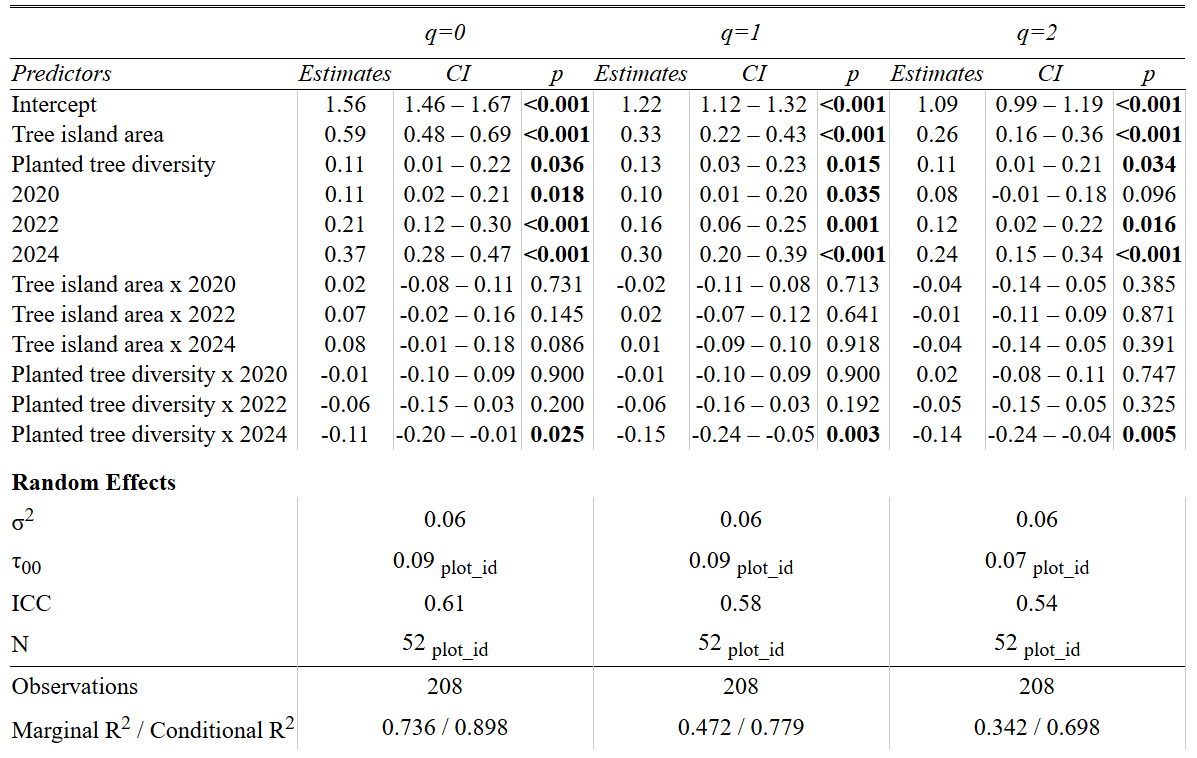

**Table S6**: LMM outputs for alien recruiting diversity at three q orders (^0^*D,* ^1^*D,* ^2^*D*).

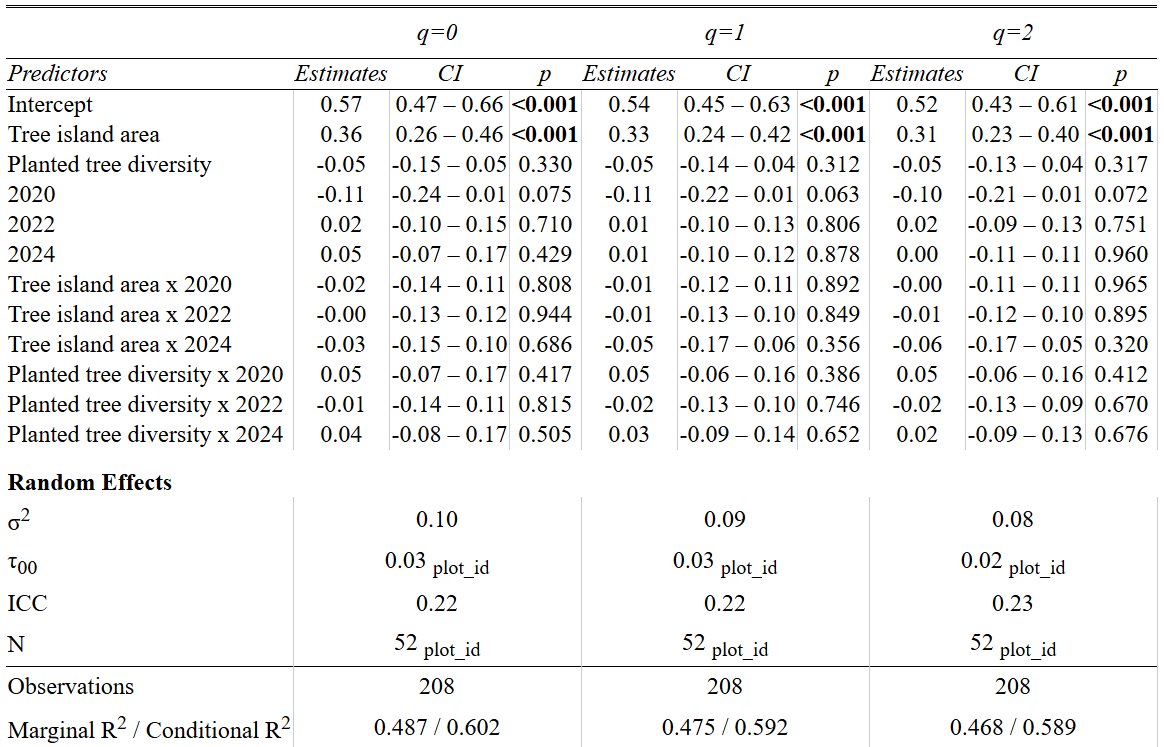

**Data processing protocol**

All data processing has been performed in R version 4.4.1 (R Core Team, 2024).

For data cleaning of the 2024 dataset, specific data entry errors were corrected manually, including corrected data about recorded individuals’ stems (mainstem vs. additional stem), species identity, tag types and tag numbers, specimen collection number, d10 values, number of stems for *Gigantochloa scortechinii* (bamboo), and information about stilt roots for *Leea javanica*. Duplicate data sheets for tree islands were removed, and zero values were changed to NAs where applicable. All family names were added corresponding to the respective species names, as well as habitat, origin, and life form information. Numbers of stems for *Gigantochloa scortechinii* (bamboo) were imputed when > 100 or when NA, and the variable was categorized into consecutive categories of 10 stems. Recorded non-woody species were removed from the data. Further, NA species were excluded from the analysis.

For cross-year data cleaning of the 2018-2024 datasets, all individual datasets for each year were harmonized and combined, allowing for adequate temporal comparisons of recruiting diversity. This included removing dead and lost stems from the datasets, as well as all additional stems, since additional stems were recorded only in 2024. Moreover, in the 2018-2024 cross-year data, all liana species were excluded because they were only recorded in 2024. After careful checks of tag numbers and exclusion of possible data entry errors, 24 stems were added to the 2022 data for which recordings in both 2018 and after 2022 were present, indicating specific data gaps for these stems only in 2022. The manually added stems were indicated in the datasets. Moreover, cross-year recorded stems detected in the field in 2024 as having vegetatively reproduced from the main stems of planted *Peronema canescens* trees were indicated in the datasets as not natural regeneration and were later excluded.

Further, reidentifications of specific individuals were aligned across all years, and new species names were adopted where species names generally changed. Unclear individual cases of *Ficus dimorpha* and *Ficus variegata* were assigned as *F. dimorpha* during data processing, as this seemed to be the much more abundant species across tree islands. For position data of stems, information on reference trees and oil palms was corrected, and x- and y-coordinates were calculated from the position data for all stems. For cases where position data for the same individual differed across years, the latest available position data was applied to all recordings for that individual across years. Further changes included removal of duplicates within one year, typos in tag numbers were corrected, old tag numbers were matched with replaced tag numbers, and all stems were given an ID base matching with the collection number of their corresponding species.
